# Global analysis of epigenetic heterogeneity identifies divergent drivers of esophageal squamous cell carcinoma

**DOI:** 10.1101/641357

**Authors:** Wei Cao, Hayan Lee, Wei Wu, Aubhishek Zaman, Sean McCorkle, Ming Yan, Justin Chen, Qinghe Xing, Nasa Sinnott-Armstrong, Hongen Xu, M.Reza Sailani, Wenxue Tang, Yuanbo Cui, Jia liu, Hongyan Guan, Pengju Lv, Xiaoyan Sun, Lei Sun, Pengli Han, Yanan Lou, Jing Chang, Jinwu Wang, Yuchi Gao, Jiancheng Guo, Gundolf Schenk, Alan Hunter Shain, Fred G. Biddle, Eric Collisson, Michael Snyder, Trever G. Bivona

**Affiliations:** Translational Medical Center, Zhengzhou Central Hospital Affiliated Zhengzhou University, Zhengzhou, China; Department of Genetics, School of Medicine, Stanford University, CA, USA; Department of Medicine, University of California San Francisco, San Francisco, CA, USA; Helen Diller Family Comprehensive Cancer Center, University of California San Francisco, San Francisco, CA, USA; Computational Science Initiative, Brookhaven National Laboratory, Upton, NY, USA; Basic Medical College, Zhengzhou University, Zhengzhou, China; Institutes of Biomedical Sciences and Children’s Hospital, Fudan University, Shanghai, China; Precision Medicine Center, The Academy of Medical Sciences, Zhengzhou University, China; Jiangsu Mai Jian Biotechnology Development Company, Wuxi, China; Department of pathology, Linzhou Cancer Hospital, Linzhou, China; Annoroad Gene Company, Beijing, China; Center for Digital Health Innovation, University of California San Francisco, San Francisco, CA, USA; Department of Dermatology, University of California San Francisco, San Francisco, CA, USA; Department of Biological Sciences, University of Calgary, Calgary, Canada

## Abstract

Epigenetic landscapes can shape physiologic and disease phenotypes. We used integrative, high resolution multi-omics methods to characterize the oncogenic drivers of esophageal squamous cell carcinoma (ESCC). We found 98% of CpGs are hypomethylated across the ESCC genome and two-thirds occur in long non-coding (lnc)RNA regions. DNA methylation and epigenetic heterogeneity both coincide with chromosomal topological alterations. Gene body methylation, polycomb repressive complex occupancy, and CTCF binding sites associate with cancer-specific gene regulation. Epigenetically-mediated activation of non-canonical WNT signaling and the lncRNA *ESCCAL-1* were validated as potential ESCC driver alterations. Gene-specific cancer driver roles of epigenetic alterations and heterogeneity are identified.

## Main Text

Epigenetic regulation is an important determinant of many biological phenotypes in both physiologic and pathophysiological contexts ^1^. However, epigenetic forces shaping the evolution of complex diseases such as cancer remain incompletely defined. Esophageal cancer is the sixth leading cause of cancer-related death and the eighth most common cancer worldwide ^2^. In China and East Asia, ESCC is the most prevalent pathohistological type of esophageal cancer ^3^. Comprehensive analysis by whole-genome and whole exome sequencing uncovered the genetic landscape of ESCC ^4–9^ and multi-region whole-exome sequencing revealed intra-tumor genetic heterogeneity in ESCC ^10^. This intra-tumor genomic heterogeneity could serve as a prognostic predictor in esophageal cancer ^11^ and as a potential foundation for improved treatment. Notable and frequently mutated epigenetic modulator genes in ESCC include *KMT2D, KMT2C, KDM6A, EP300* and *CREBBP*, and epigenetic perturbations might interact with other somatic genomic alterations to promote ESCC. The interplay between epigenetic perturbations and other somatic genetic alterations may have a critical role during ESCC tumorigenesis ^4^.

The Cancer Genome Atlas research group (TCGA) identified ESCC-related biomarkers at a multi-omics level (genomic, epigenomic, transcriptomic, and proteomic) and pinpointed 82 altered DNA methylation events, along with altered transcriptional targets genomic alterations ^9^. While genomic and transcriptomic-level studies of ESCC produced valuable biological discoveries and resources, the single-nucleotide resolution of the epigenetic landscape of ESCC, and of most other cancers, at the whole genome level remains poorly studied. This knowledge gap is due to the comparatively high cost, computational complexity, and technical challenges of capturing genome-wide and single-nucleotide resolution of the epigenetic landscape. Consequently, an integrative and causal analysis across orthogonal multi-omics datasets remains incomplete.

We addressed this challenge by using an integrated multi-omics study that includes whole genome bisulfite sequencing (WGBS), whole genome sequencing (WGS), whole transcriptome sequencing (RNA-seq) and proteomic experiments on a cohort of ESCC samples and their adjacent non-tumor esophageal tissues along with orthogonal analysis and validation using the large TCGA-esophageal cancer (ESCA) dataset. Our goal was to understand the extent and complexity of epigenetic heterogeneity in DNA methylation alterations and consequent dysregulation of both protein coding and non-coding gene expression.

## Results

### Whole genome bisulfite sequencing reveals the epigenetic landscape and heterogeneity in ESCC

Different types of cancers exhibit unique epigenetic alterations, particularly in the DNA methylome ^12–14^. We initially collected ten pairs of primary ESCC samples and their adjacent non-tumor tissues (Supplementary Fig.1), performed WGBS with over 99% of a bisulfite conversion ratio, and generated a mean 15x sequencing depth per sample (Supplementary Table 1, Supplementary Fig. 2). Over 99% of CpG dinucleotides were covered and ∼95% of CpGs were reliably mapped by more than five reads. To ensure the quality of data, bisulfite converted sequencing reads were aligned with TCGA-ESCA Human Methylation 450K (HM450K) array (Supplementary Fig. 3a) and showed strong concordance in all normal and tumor samples (Pearson r = 0.9644, p-value < 0.01, Supplementary Fig. 3b); the coefficient for WGBS-ESCC versus TCGA-ESCC, or TCGA-EAC (esophageal adenocarcinoma) was 0.7570 and 0.5554, respectively (Supplementary Fig. 3c, 3d). DNA methylation at non-CpG contexts was present in less than 0.5% in our samples.

More than 5 million differentially methylated cytosines (DMCs) were identified using a one-way ANOVA test (FDR < 5%) (Fig. 1a). Among them, 57.5% were located at known annotated regions, 42.5% were located at unannotated regions of the genome (Supplementary Fig. 4a). Methylation loss in cytosines in ESCC accounted for 97.3% of the DMCs and was mostly confined to intergenic regions of the genome. Only 2.7% of the DMCs were gains of methylation in ESCC (proportional test for hyper- and hypo-methylation, p value < 2.2e-16, Fig. 1b) and 83.67% of them mapped to gene bodies, promoters, and enhancers with RefSeq annotation (Supplementary Fig. 4b). Of the hypomethylated DMCs in ESCC, 63.08% were mapped to lncRNA regions with ENCODE annotation (v27lift37), whereas 58.01% of hypermethylated DMCs in ESCC were dispersed in antisense RNA regions of the genome (Supplementary Fig. 5a and 5b). These DMCs clearly discriminate normal tissues from tumor tissues, as measured by unsupervised Principal Component Analysis (PCA) (Fig. 1c), similar to unsupervised transcriptome-mediated clustering of normal and tumor samples (PCA in Supplementary Fig. 6a and Dendrogram in 6b). In the larger sample set (n=202) of TCGA-ESCA, the differentially methylated CpG probes present in the lower resolution Illumina HM450K array were also able to discriminate normal and tumor and even subtypes of esophageal cancer using t-distributed stochastic neighbor embedding (t-SNE), a nonlinear dimensionality reduction algorithm (Fig. 1d). The data suggest that alterations in DNA methylation can characterize the biological features of cellular states of physiologic and pathophysiologic phenotypes.

**Figure 1.**
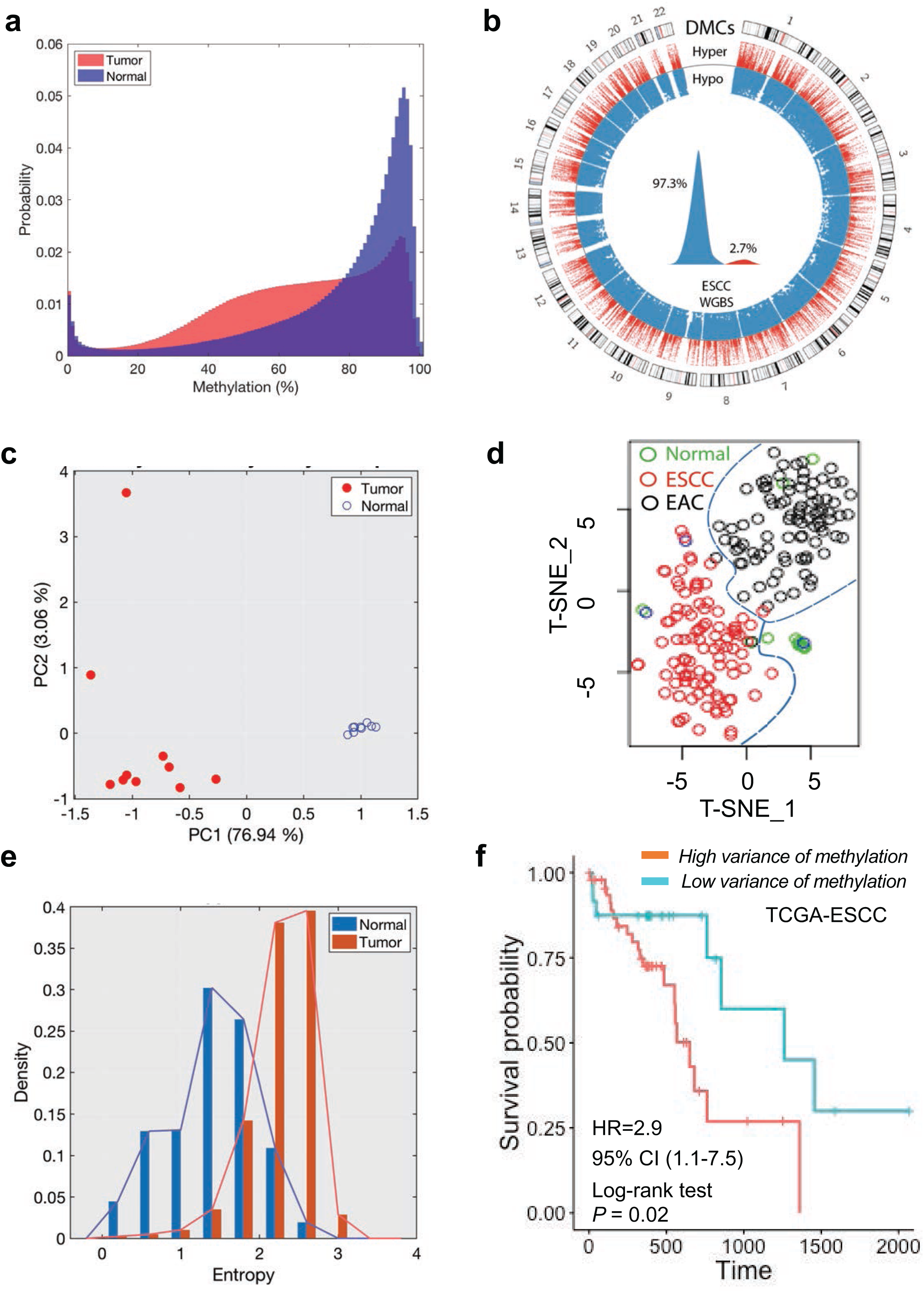
Whole genome bisulfite sequencing (WGBS) unveiled global hypomethylation and heterogeneity in esophageal squamous cell carcinoma (ESCC). (a) The asymmetric density distribution of all CpG methylation statuses in the normal esophageal tissues versus ESCC. ESCCs lose methylation which leaves most CpGs partially methylated. (b) Circos plot of 5 million differentially methylated CpGs (DMCs) between ESCC tumor and adjacent normal tissue. DMCs are substantially hypomethylated in ESCC (97.3%). Only 2.7% are hypermethylated in ESCC. (c) Principal Component Analysis (PCA) shows DMCs discriminate tumor samples from normal samples. (d) t-Distributed Stochastic Neighbor Embedding (t-SNE) showed CpG methylation profiling of TCGA-esophageal cancer from human methylation 450K analysis clustered into either normal tissue (green circles) or ESCC (red circles) or esophageal adenocarcinoma (black circles) subtypes. (e) Entropy analysis of all CpGs showed variations per CpG in normal esophageal tissues (blue bars) and ESCC (red bars). The entropy of CpGs in ESCC was higher than in normal samples. (f) Kaplan-Meier survival analysis demonstrated TCGA-ESCC patients (N=73) with higher variance (the sum of squared distance of each CpG methylation from the mean) of CpG methylation in tumors showed worse survival time than those with lower variance. Log-rank p-value=0.02.

DNA methylation heterogeneity has been observed in other cancer types ^14,15^ and stochastically increasing variation in DNA methylation appears to be a property of the cancer epigenome ^16^. The clinical significance of such inferences remains unclear. We found a higher variance of altered methylation in ESCC (275.76 ± 204.01) compared with normal samples (95.67 ± 112.38, two sample t-test p-value ≈ 0) in our cohort as well as in the TCGA-ESCC cohort (p < 2.2e-16) (Supplementary Fig. 6c). As a further measurement of the level of epigenetic variance, we calculated Shannon’s entropy of methylation levels at each CpG locus. We observed increased entropy in ESCC compared with normal samples (two sample t-test p-value ≈ 0) (Fig.1e), and this is consistent with the increase in stochastic ‘noise’ (heterogeneity) in tumors. Our simulation using the Euler-Murayama method ^17^ also reflected increased DNA methylation heterogeneity in ESCC (Supplementary Fig. 6d). Using the independent TCGA-ESCC clinical cohort, we stratified patient samples by their variance of methylation level and found that the group with a higher variance (N=49) of methylation levels showed a worse overall survival time (hazard ratio=2.9, 95% confidence interval (1.1∼ 7.5), p-value < 0.05) (Fig. 1f). This provides potential clinical relevance of the epigenetic heterogeneity that we uncovered in ESCC.

### Differentially methylated regions (DMRs) associate with alterations of genome topology and a global abnormal functional annotation of the ESCC transcriptome

We further defined 299,703 DMRs (p value <= 0.05, FDR < = 0.05) between tumor and normal tissues, resulting from a CpG density peak of 4% and a DMR peak size of 200-400 base pairs (bp) (Fig. 2a, Supplementary Fig. 7a and 7b). Only 1.8 % of these DMRs are hypermethylated, while 98.2 % of DMRs are hypomethylated (proportional test, p-value < 2.2e-16) in tumors relative to normal tissues. DMRs in regions of −2990bp ∼ +6990bp appear hyper-methylated while gene bodies, intergenic, and non-coding regions are in general hypomethylated in tumors (Fig. 2b, Supplementary Fig. 7c,7d). The occupancy of each transcription factor (TF) binding consensus varies in the genome (161 TF binding sites from ENCODE), with POLR2A (5.23%) and CTCF (3.55%) ranking at the top (Supplementary Fig. 8a, Supplementary table 2). We searched CpG content in these TF binding sequence and the top 20 TFs affected by methylation alterations in consensus binding sites were identified. Notably, the Polycomb Repressor Complex 2 (PRC2) subunits SUZ12 and EZH2 binding sites were substantially affected by hypermethylation in the CpGs (Supplementary Fig. 8b-8d). These observations indicated the possibility of a paradoxical activation mechanism for PRC2 target genes through loss of PRC2 occupancy in gene promoters in tumor cells.

**Figure 2.**
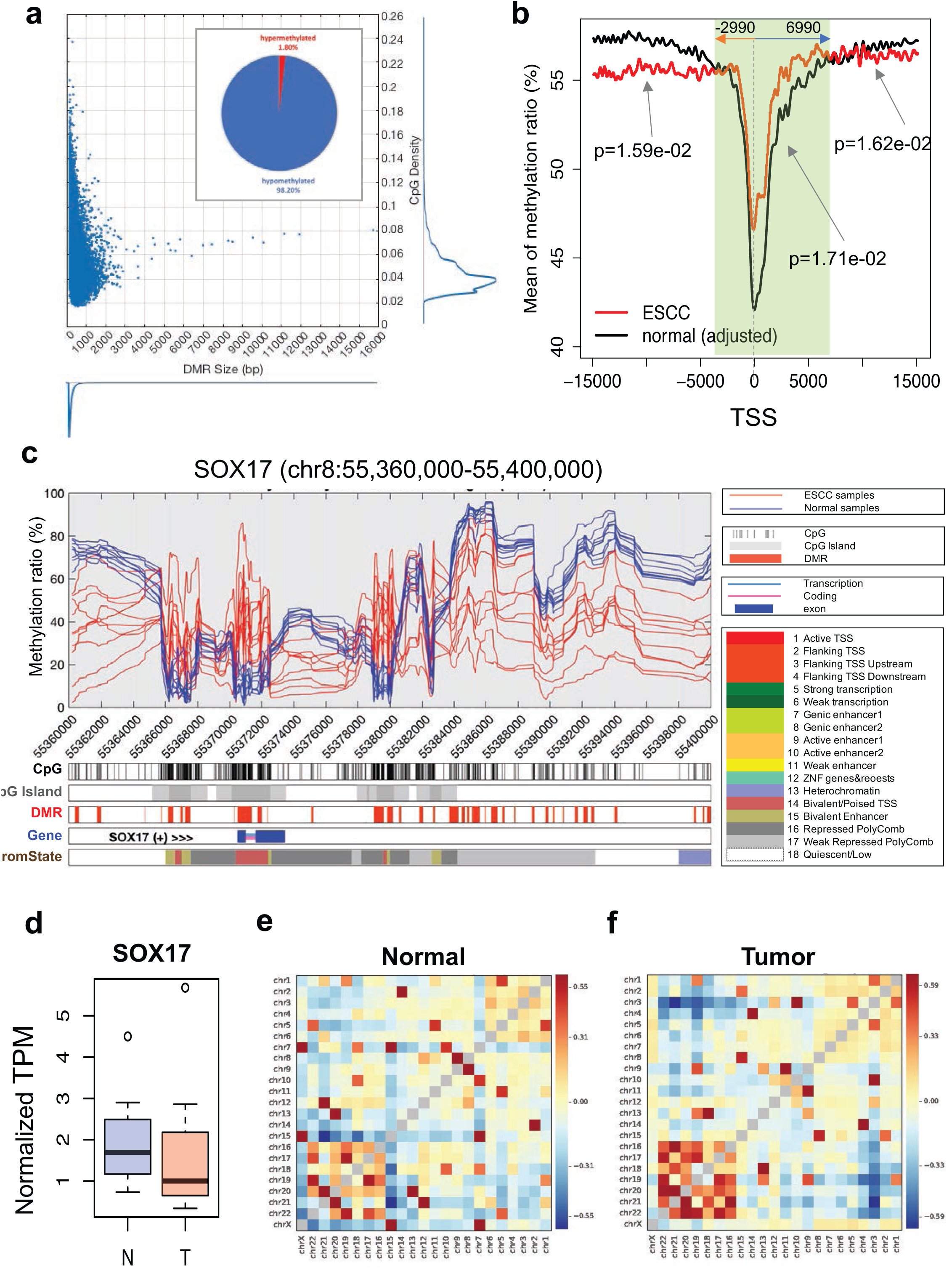
Differentially methylated regions (DMRs) and their functional impacts on the ESCC genome. (a) A DMR identification algorithm from DMC was developed using two criteria: (1) two flanking DMCs should be close (150 base pairs, bp) given the minimum size of CGI 150bp; and (2) the methylation pattern should be consistent, either hypomethylated or hypermethylated within a DMR. Our algorithm revealed the landscape of DMRs in terms of DMR size and CpG density. Both distributions of DMR size and CpG density are asymmetric and have long tails as DMR size increases in length and CpG density is more compact. The peak of DMR size is 200-300bp and the peak of CpG density is approximately 4%. (b) Methylation level of CpGs within 15,000 bp upstream and downstream relative to a TSS were assessed in ESCC and normal esophagus separately. Overall methylation conversions between normal esophagus tissue and ESCC were observed. CpGs in ESCC tend to be hypermethylated between 3,000 bp upstream and 7,000 bp downstream of a given TSS. The arrow indicates the p value for the specific region of significant methylation changes. (c) A representative genomic region at chr8:55,360,000-55,400,000 with hypermethylation in CpG island and hypomethylation in the CpG shore. (d) SOX17 expression is decreased in ESCC tumors (T) relative to normal tissue (N). (e-f) The chromosomal interaction changes during tumorigenesis as detected by genome-wide chromosome conformational capture assay (Hi-C). There are closer interactions among chr16 through chr22 in tumor compared to normal tissue.

The DMRs were distributed mostly (>6%) in Chromosome (Chr)8, Chr19 and Chr20 after normalization to chromosome size (Supplementary Fig. 9a) and DMRs are enriched (>20%) in gene promoters at Chr19 (Supplementary Fig. 9b). We integrated the most significant DMRs with all CpGs, CpG island, chromatin state, and potential TF binding data using the ENCODE dataset ^18^. We observed Chr8 harbors three large genomic regions with unique DMR patterns and these regions contain the *SOX17, RGS22*, and *ESCCAL-1* (*CASC9*) gene loci, respectively. For example, around the gene of *SOX17* (Chr8:55,360,000-55,400,000), CpG island regions were hyper-methylated but CpG shore regions were hypo-methylated (Fig. 2c). Two CpG islands with significant hyper-methylation upstream of the *SOX17* gene were observed but there is no association with low gene expression (p-value = 0.668, Fig. 2d). In the region 100,650,000-101,190,000) of Chr8, hypo-DMRs covered all of the gene body of *RGS22* (regulator of G protein signaling) (Supplementary Fig. 10), which is a putative tumor suppressor ^19^. The region around the lncRNA ESCCAL-1 (Chr8:76,130,000-76,240,000), which was previously identified by us ^20^, contained significantly hypo-methylated DMRs in its promoters and we further investigated the uncharacterized biological function of this lncRNA later in this study.

A link between hypo-methylated blocks, variable gene expression, and large heterochromatin domains such as Large Organized Chromatin lysine (“K”) modification (LOCK) or lamina-associated domains (LAD) was previously reported in cancer ^21^. Nevertheless, the relation between significant DMRs and Topologically Associating Domains (TADs), or self-interacting genomic regions ^22^ is largely uncharacterized in ESCC. Genome-wide chromosome conformation capture followed by massively parallel DNA sequencing (Hi-C) showed increased TAD abundance and reduced TAD size in the ESCC cancer genome relative to the normal genome (Supplementary Fig.11a). The interactions between chromosomes was altered during ESCC tumorigenesis (Supplementary Fig.11b). Closer interactions between Chr16 through Chr22 were observed in ESCC compared with normal esophageal cells (Fig. 2e and 2f). Using the Hi-C data, we inferred two compartments: open euchromatin of transcriptionally active states (compartment A) and closed chromatin of transcriptionally silent states (compartment B) ^22^. We observed 22.61% of the compartment shift (A→B or B→A) during ESCC tumorigenesis (Supplementary Fig. 11c). The A→ B shifted regions contain 0.5∼9% DMRs with Chr10 and Chr19 showing the most DMR occupancy (Supplementary Fig.12a-d). In contrast, the B→A shifted regions have 0.5∼20% DMRs with a higher percentage of DMR in Chr3 (Supplementary Fig.13a-d). This suggests that a functional link exists in ESCC between DMR alterations and shifts in genomic architecture.

Significant methylation changes were identified by WGBS at 5085 gene promoter regions (−4500bp ∼500bp relative to a TSS). Gene set enrichment analysis (GSEA) analysis of these target genes harboring promoter hypomethylation indicated an over-representation of WNT/β-catenin signaling, whereas gene promoters harboring hyper-methylation were enriched for KIT signaling genes (Supplementary Fig. 14a). In parallel, GSEA analysis for differentially expressed genes (DEGs) from the RNAseq dataset indicated enrichment for genes regulating cell cycle pathways and metallopeptidase activity (Supplementary Fig. 14b). Hence, the DMRs associated with alterations of genome architecture appear to shift the gene regulatory networks during ESCC tumorigenesis.

### Aberrant DNA methylation in promoter regions mediates transcriptional dysregulation in ESCC

DNA methylation at regulatory regions influence transcript expression levels ^23^. From our WGBS analysis, we identified 5085 promoter regions (−4500bp ∼ +500 bp to TSS) of coding and non-coding genes whose CpGs were significantly differentially methylated (FDR < 0.01) and focused on the 4768 significantly differentially expressed transcripts in ESCC relative to the adjacent normal tissues from the RNA-seq dataset. We then identified 694 genes that showed significant differential methylation alteration in promoters and concomitant dysregulation of gene expression (Fig. 3a). The genes were systematically classified into four distinct clusters (denoted as C1, C2, C3 and C4) according to methylation (met) and gene expression (ge) pattern. C1(H_met_L_ge_) showed hypermethylated promoters with decreased ge; C2(L_met_H_ge_) showed hypomethylated promoters with increased ge; C3(H_met_H_ge_) contained hypermethylated promoters with increased ge; C4(L_met_L_ge_) denoted hypomethylated promoters with decreased ge. C1 and C2 followed the well-documented canonical model, showing anti-correlation in promotor methylation and gene expression ^13^; in contrast, genes in C3 and C4 showed a non-canonical pattern in that promotor methylation and gene expression were positively correlated (Fig. 3b, 3c, Supplementary Fig. 15, Supplementary table 3). Among the 694 genes, only 1.5% of them harbor non-synonymous mutations from our selected cases of WGS (Supplementary Fig. 16) and no copy number changes of these genes were inferred from RNA-seq (Supplementary Fig. 17). Therefore, the majority (98.5%) of these dysregulated genes in cluster C1∼C4 appear to occur via epigenetic dysregulation (epimutation) ^24^. This phenomenon is also seen in the independent TCGA-ESCC (n=96) sample cohort, by analysis of the available multi-OMICs dataset (Supplementary Fig. 18a-d).

**Figure 3.**
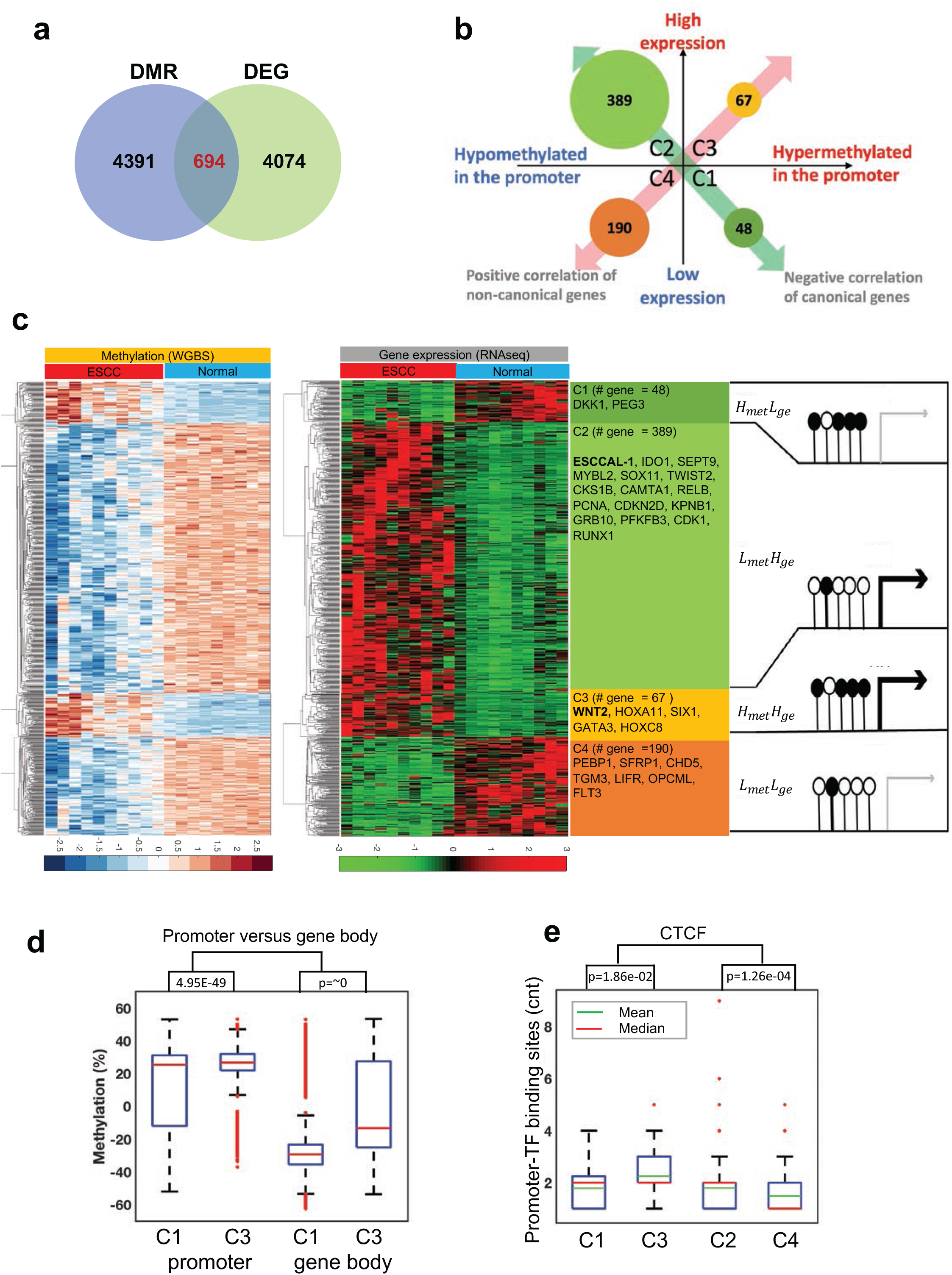
Integrative analysis of WGBS and RNAseq uncovered methylation-mediated diverse gene regulation based on promoter methylation levels. (a) A total of 694 genes were selected for the methylome-transcriptome association analysis. The selected genes have statistically significant DMRs (FDR <= 0.001) in promoters, defined as 4500 base pair (bp) upstream and 500bp downstream relative to transcription start sites, and are statistically significant DEGs (Differentially Expressed Genes) (FDR <= 0.05). (b-c) Further association of promoter methylation and expression of the 694 genes identified. There are four clusters: C1: Genes that are hypermethylated in the promoter with low expression level in ESCC; C2: Genes that are hypomethylated in the promoter with high expression in ESCC; C3: Genes that are hypermethylated in the promoter with high expression in ESCC; C4: Genes that are hypomethylated in the promoter with low expression in ESCC. Genes in C1 and C2 fit the canonical model of regulation, while genes in C3 and C4 are not well explained by current understanding. Representative genes are listed in each cluster. (d) The quantification of CpG methylation in gene promoters and gene bodies in C3 is significantly higher than in C1, p < 0.001. (e) CTCF binding sites are significantly higher in C3, indicating hypermethylation of inhibitors leads to de-repression to promote gene expression.

The underlying mechanisms of the divergent regulation of gene expression are complex and involve DNA methylation, chromatin remodeling, and DNA accessibility ^25^. We explored a potential explanation for the non-canonical patterns in C3 and C4 across epigenetic regulatory features. First, we cross-referenced 13,000 TCGA-ESCA chromatin accessible regions as determined by ATAC-seq ^25^ to the regulatory regions of the 694 genes. Accessibility in promoter regions was highly associated with gene expression in C2 and C3 (Supplementary Fig. 19a, b). Second, methylation in defined promoter regions and in gene bodies showed a differential phenotype between C1 and C3 (but not between C2 and C4): methylation levels in gene bodies were higher in C3 (−5.4587 ± 26.3450) than C1 (−26.8551 ± 16.4716, p < 0.05) (Fig. 3d). Third, hyper-methylation at cohesion and CCCTC-binding factor (CTCF) binding sites could compromise binding of this methylation-sensitive insulator protein and result in gene activation ^26^. Thus, we searched for CTCF binding sites within promoter regions of the 694 genes and observed that the CTCF binding sites were enriched in C3 (Fig. 3e), which could partially explain the phenotype of high promoter methylation and high gene expression. Fourth, the compartment shift regions inferred from Hi-C data showed that 53.24% of the genes in C3 shifted from a closed state to an active state (Supplementary Fig. 20). The data also indicated that the promoters of genes in C4, despite being hypomethylated in the tumor, were inaccessible. This highlights both impotence of accessibility and absence of methylation as linked features of the gene expression pattern in the C4 cohort. Gene enrichment analysis of the 694 genes was performed using multiple databases (KEGG ^27^, WikiPathways ^28^, ENCODE ^18^, ChEA^29^) and showed PRC2 subunits (EZH2 and SUZ12)-mediated polycomb repressive gene sets were enriched in the non-canonical clusters C3 and C4 (Fig. 4a). We searched for ENCODE-defined EZH2 and SUZ12 binding sites across gene promoters in C1-C4 and observed that EZH2 occupancy was enriched in C3 (1.5970 ± 1.2316) and C4 (0.6000 ± 0.7684) compared with C1(0.9167 ± 0.8464) and C2 (0.2336 ± 0.5870), respectively (p-value < 0.001) (Fig. 4b). SUZ12 occupancy was higher only in C3 gene promoter regions (1.5522 ± 1.7946), (Fig. 4c). To understand the functional mechanism that is responsible for differential methylation at target gene promoters, we performed unsupervised hierarchical clustering of ENCODE-defined known TF binding sites in the DMRs of the 694 gene promoters. The analysis showed that EZH2 binding sites were enriched in genes in C3 compared to other clusters (Fig. 4d, Supplementary Fig. 21). In addition, we performed a correlation analysis between gene expression difference (expression fold change) and corresponding promoter methylation level difference (methylation Δ) of genes in C3. WNT2 was identified as the top non-canonical gene in C3 (Fig. 4e). The collective data show the non-canonical gene expression pattern (C3) appears to arise via de-repression of the EZH2-mediated suppressor effects on promoter regions of genes in C3 to increase gene expression, which we experimentally validate later.

**Figure 4.**
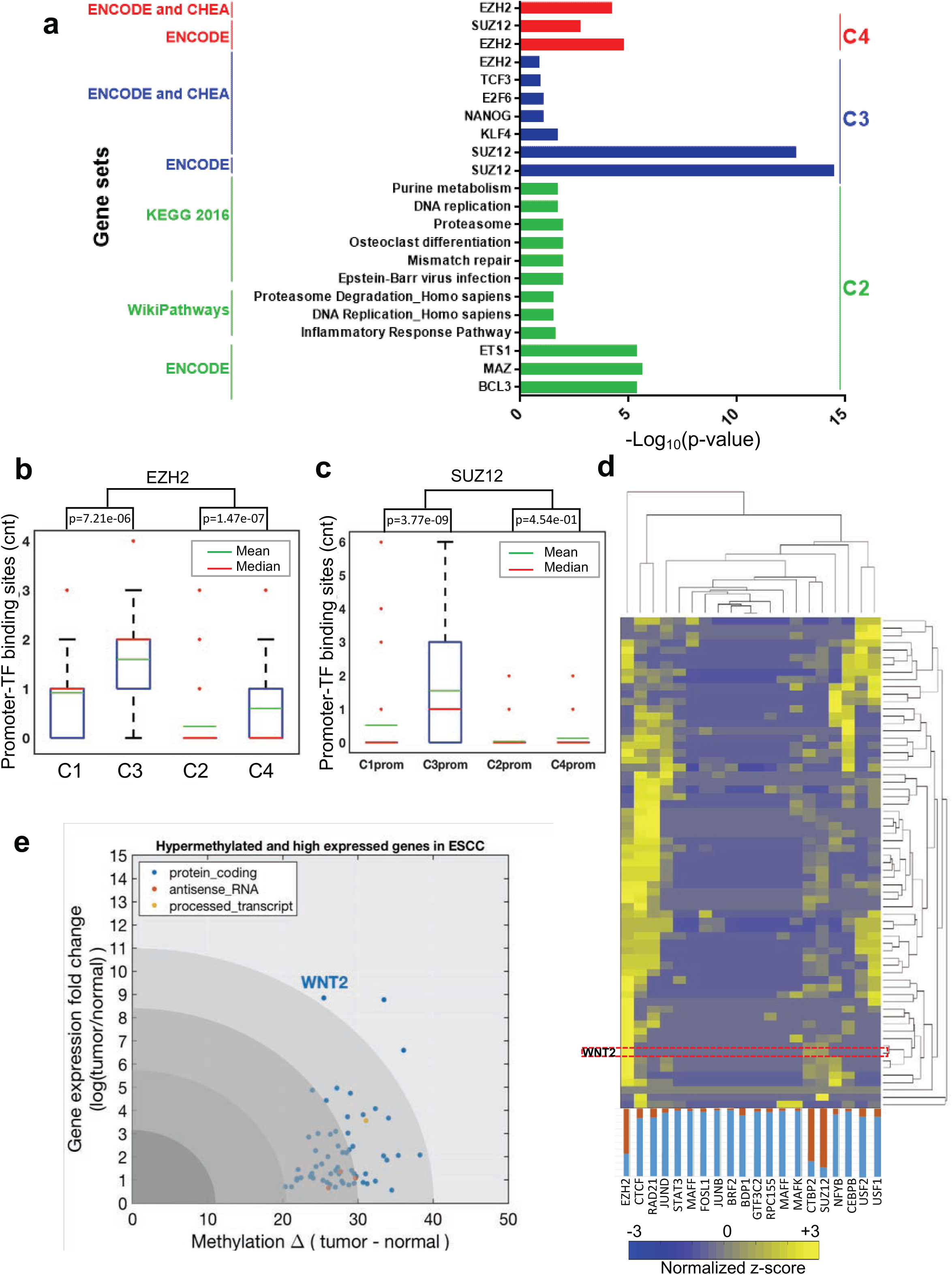
DNA methylation at regulatory protein consensus protein binding sites and impacts on gene expression. (a) Gene Set Enrichment Analysis (GSEA) analyses for the four distinct methylation-transcriptome clusters. Lists of genes in C2, C3 and C4 were subjected to GSEA using hypergeometric statistics for gene sets collected from multiple databases (ENCODE, CHEA, KEGG, WikiPathways, Reactome, GO molecular function, Panther, BIOGRID etc.). The significance of the hypergeometric analysis is indicated as –Log_10_ (p-value) in the form of a horizontal histogram where bar heights represent level of significance. Bars are color coded based on their inclusion in each cluster. GSEA revealed gene pathways or GO terms in different clusters, including polycomb repression complex2 (PRC2) subunit, EZH2 (Ester of Zinc Finger Homolog 2), and SUZ12 (Polycomb Repressive Complex 2 Subunit) binding sites significantly enriched in C3, suggesting hypermethylation of PRC2 de-represses gene expression. Genes in C1 have no significant gene set enrichment. (b-c) In silico analysis of EZH2 and SUZ12 binding sites in each cluster and gene promoters show more significant binding scores in C3 than in other groups, p-value < 0.001. (d) The probability of the top 20 transcription factor consensus binding sites in C3 showed EZH2 has the highest binding scores in a subset of genes in C3, including WNT2. The heatmaps for C1, C2, and C4 are in Supplementary Figure 21. (e) In non-canonical gene cluster C3, WNT2 is significantly hypermethylated in the promoter (FDR=6.6005e-03) and highly expressed in ESCC (FDR=0.0039). WNT2 also shows the highest fold change in gene expression in ESCC relative to adjacent normal tissues.

### DNA methylation gain at the promoter region activates WNT2/β-catenin pathway in ESCC

Epigenetics dysregulation of the components of MAPK, AKT and WNT pathway can promote aberrant activation of these pro-growth pathways in ESCC ^30^. We extracted known components of these genes from published literature ^30,31,32^ and compared their gene expressions between tumor and normal samples. The gene expression analyses of the component genes in MAPK, AKT and WNT pathway identified only WNT2 in WNT pathway was significantly highly expressed in the tumor samples compared to normal samples (Supplementary Fig. 22a, b). These data indicate selective and specific upregulation WNT2 in ESCC tumors through a putative non-canonical epigenetic regulatory mechanism. WNT2 belongs to the structurally related WNT family of genes that functions as secretory ligands for the WNT signaling pathway ^33^. Canonical WNT signaling pathway results in stabilization of the transcriptional co-regulator β-catenin and subsequent upregulation of downstream target genes ^33^.

To gain mechanistic insight into the epigenetic regulation of *WNT2* promoter, we queried our transcription-factor target gene hierarchical clustering analysis for genes in C3 and found that the EZH2 binding site, along with SUZ12 binding sites, are present at the *WNT2* promoter region compared to other transcription factors (Fig. 4d). EZH2 and SUZ12 are subunits of PRC2, which has histone methyltransferase activity to primarily tri-methylate histone H3 on lysine 27 (H3K27me3) ^34^. We also found that the EZH2 biding site at *WNT2* promoter overlap with the hyper CpG methylation sites at *WNT2* promoter region in cancer cells (Figure 5a).

**Figure 5.**
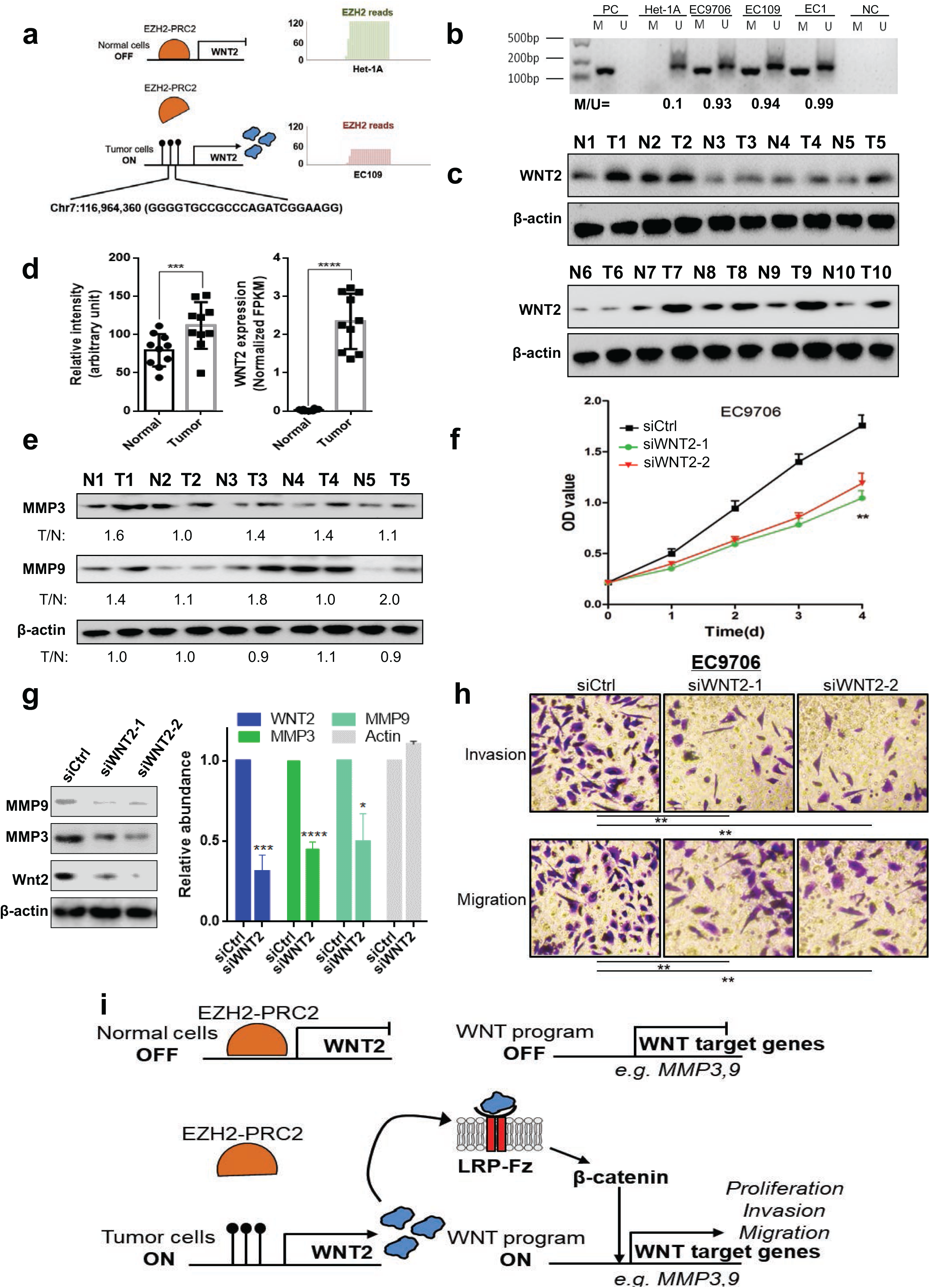
Non-canonical hypermethylation in the WNT2 promoter leads to high WNT2 expression in ESCC. (a) Chromatin immunoprecipitation by EZH2 protein binding followed by high-throughput DNA sequencing (ChIP-seq) showed enriched reads on WNT2 promoter region in normal cells, but not in ESCC cells. (b) The WNT2 promoter region is hypermethylated in ESCC cell lines by methylation specific PCR analysis. M: methylation detection. U: unmethylation detection. PC: positive control. NC: negative control. HET-1 cells are an immortalized normal esophageal epithelial cell line. EC9706, EC109 and EC1 are patient-derived ESCC cell lines. (c) WNT2 is overexpressed in the tumor samples. Ten paired tumor and normal tissue samples were collected and lysed for western blot analysis for the indicated proteins using SDS-PAGE. (d) Band intensity quantification for the western blot results are indicated on the left as a bar diagram. On the right, normalized fragment per kilobase of transcript (FPKM) value indicative of the RNA abundance for 10 pairs of normal and tumor samples are indicated as a bar graph. Significance for comparison between the two cohorts was measured using an unpaired student’s t-test (p-value < 0.01). (e) MMP3 and MMP9 are highly expressed in the tumors. Five paired tumor and normal tissue samples were collected and lysed for western blot analysis for MMP3, MMP9 expression. (f) WNT2 depletion with two independent siRNAs inhibits growth in the ESCC cell line EC9706. The MTT assay was performed in siControl and WNT2-silenced samples and represented as a line plot showing normalized optical density (OD) values as a representation of cell numbers at serial time points. Significance between the two cohorts was measured using a paired student’s t-test (p-value < 0.01). Data are representative of three independent experiments. (g) Two independent siRNAs for WNT2 knockdown decrease MMP3 and MMP9 expression. Cells were harvested from control and WNT2-depleted cells and expression of the indicated proteins was measured using SDS-PAGE and Western blot. Data are representative of three replicates. Significance for band intensity comparison between the siControl and WNT2 knockdown cohorts was measured using an unpaired student’s t-test (p-value < 0.01). (h) Two independent siRNAs to silence WNT2 expression reduced the invasion and migration of ESCC cells relative to siRNA controls. p-value < 0.001. (i) Schematic representation of the mechanism of EZH2/PRC2-WNT2-MMP signaling upregulation in ESCC.

*WNT2* promoter region (Chr7: 116,960,000-116,965,000) was hypermethylated but paradoxically WNT2 gene expression was increased in tumors (Supplementary Fig. 23 from our WGBS dataset, Supplementary Fig. 24 from TCGA-dataset). We reasoned that de-repression of EZH2 occupancy may cause non-canonical methylation-mediated activation of WNT2 gene expression in ESCC, we validated EZH2 occupancy on the differentially methylated regions of *WNT2* promoter by performing Chromatin ImmunoPrecipitation sequencing (ChIP-seq) in normal immortalized esophageal epithelial cells (Het-1A) and the patient-derived ESCC cell line, EC109. The ChIP-seq analysis showed EZH2 binding peaks at *WNT2* promoter region in normal cells compared to minimal binding peaks in the ESCC cells (Fig. 5a). Furthermore, we confirmed the promoter region of *WNT2* was hypermethylated in three esophageal cancer cell lines (EC9706, EC109, and EC1) while no methylation was detected in normal esophagus epithelial cells (Het-1A) (Fig. 5b). In addition, WNT2 mRNA and protein expression was also higher in independent ESCC samples (Fig. 5c, 5d).

To identify downstream effector genes of WNT/β-catenin signaling that might promote ESCC, the GSEA of differential expression from proteomic data and RNAseq data found extracellular matrix organization (Supplementary Fig. 25) and extra-cellular metalloproteins MMP3 and MMP9 (known β-catenin targets) ^35^ gene set (Supplementary Fig. 26) are enriched in tumor samples. We validated that both MMP3 and MMP9 transcripts and proteins were highly expressed in ESCC relative to normal tissues (Fig. 5e). To test whether WNT2-mediated signaling was required for tumor cell growth, we suppressed WNT2 expression using two independent short interfering (si)RNAs in two different patient-derived ESCC cell lines (EC9706 and EC109). WNT2 knockdown significantly inhibited ESCC cell growth (p-value < 0.01) (Fig. 5f). These data place WNT2 as an essential gene for ESCC cancer cell growth. Furthermore, since MMPs are known downstream targets of WNT/β-catenin signaling activation, we tested the effect of WNT2 knockdown on MMP3 and MMP9 expression. We found baseline WNT2 expression in ESCC cell lines to be significantly higher than normal cell lines (Supplementary Fig. 27a). Furthermore, WNT2 knockdown decreased MMP3 and MMP9 expression (Fig. 5g). Since matrix metalloproteinases (MMPs) can promote tumor invasion and metastasis ^36^, we next tested whether WNT2 knockdown abrogates the migratory and invasive potential of ESCC tumor cells. In two patient-derived ESCC cell lines (EC9706 and EC109), silencing of WNT2 significantly reduced cellular invasion and migration (Fig. 5h, p-value < 0.01, Supplementary Fig. 27b, c, d, e, f). We also found that β-catenin protein expression encoded by *CTNNB1* gene is significantly higher in ESCC tumors (Supplementary Fig. 27g). Knockdown of WNT2 remarkably reduced protein level of β-catenin (Supplementary Fig. 27h). These data show that a WNT2/β-catenin/MMP3/9 signaling axis was not only required for tumor cell growth, but also for tumor cell migration and invasion in ESCC. Taken together, our data demonstrate a novel non-canonical mechanism for increased WNT2 expression in the absence of EZH2-PRC2 occupancy of the WNT2 promoter with hypermethylated CpG. The data delineate specific downstream targets of WNT2-mediated signaling and their functional consequences in ESCC (Fig. 5i).

### Epigenetic activation of the lncRNA ESCCAL-1 is a novel ESCC cancer driver gene

Increasing evidence indicates dysregulation of lncRNAs during cancer progression and metastasis, but the mechanisms of dysregulation and of action of lncRNAs in cancer are relatively poorly understood ^37^. Our WGBS analysis revealed that hypomethylated DMCs are significantly enriched in genomic regions harboring annotated lncRNAs (Fig. 6a). In the canonical cluster C2, gene expression was strongly anti-correlated with gene promoter CpG methylation levels (Fig. 6b). Previously, we showed that the lncRNA ESCCAL-1 was overexpressed in ESCC ^20^, and overexpression of its ESCCAL-1 has been reported in other cancer types ^38–41^. The mechanism underling ESCCAL-1 upregulation in cancer is unknown. We found ESCCAL-1 is one of the most notable candidates for DNA methylation loss-mediated increased gene expression in C2 (Fig. 6b). One DMR in the promoter of ESCCAL-1 showed decreased CpG methylation in cancer, leading to increased transcription of lncRNA ESCCAL-1 (Fig. 6c) ^42^. There is no mutation or copy number variation of ESCCAL-1 reported or observed in TCGA-ESCA genomic dataset or in our WGS data of three ESCC patients (Fig. 6c, top panel). Independent verification of the methylation status of the ESCCAL-1 promoter region showed 62.5% (20/32) hypomethylation in ESCC tumors versus 71.8% (23/32) hypermethylation in adjacent normal tissues, chi square test p-value < 0.01 (Fig.6e, f). In agreement, ESCCAL-1 expression was significantly higher in ESCC compared to adjacent normal tissues (p-value = 0.00113, FDR <0.05) (Fig. 6d from RNAseq). We corroborated these observations by analysis of an independent cohort of 73 ESCC tissues relative to their normal counterparts (Fig. 6g). We also noted a hypermethylated ESCCAL-1 promoter region in normal esophageal cells (Het-1A), whereas methylation was not detected in three ESCC cell lines (EC1, EC109 and EC9706) (Fig. 7a). ESCCAL-1 expression was substantially overexpressed in ESCC cell lines compared with normal cells Het-1A (Fig. 7b). Furthermore, increased expression of ESCCAL-1 was a biomarker of worse overall survival time and progression-free survival time in ESCC patients (Fig. 7c, d). Knockdown of ESCCAL-1 reduced growth of patient-derived ESCC cells *in vitro* (Supplementary Fig. 28) and *in vivo* (Fig. 7e, f).

**Figure 6.**
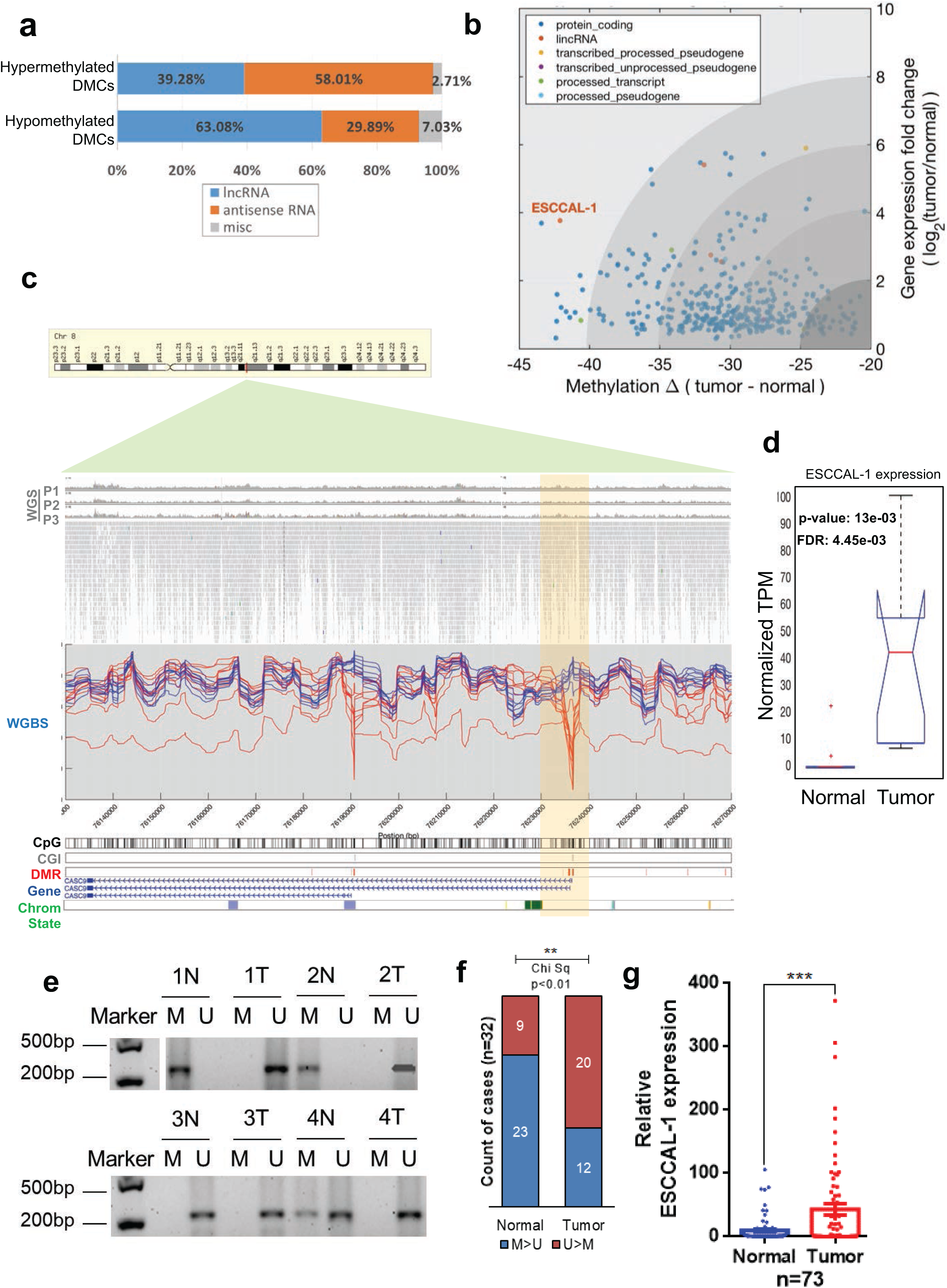
Hypomethylation-mediated upregulation of long non-coding RNAs (lncRNAs) in ESCC. (a) DMCs are mapped to functional annotated regions. 58.01% of hypermethylated DMCs overlap with antisense RNA (asRNA), whereas 63.08% of hypomethylated DMCs overlap with lncRNAs. (b) In canonical gene cluster C2, ESCCAL-1 is significantly hypomethylated in the promoter (FDR=1.7386e-04) and highly expressed in ESCC (FDR=0.01). ESCCAL-1 also shows the most significant and substantial methylation difference between normal esophageal tissues and ESCC among the lncRNAs in C2. (c) Loss of CpG methylation at the ESCCAL-1 promoter region in ESCC (chr8:76,135,639-76,236,976 of GRCh37/hg19). WGS (Whole Genome Sequencing) of three ESCC patients shows no mutation or copy number variations detected at the above indicated region. This is validated by TCGA ESCA data (N=186), where no mutation or copy number variations were observed (http://firebrowse.org/?cohort=ESCA). Whole Genome Bisulfite Sequencing (WGBS) reveals tens of DMRs around ESCCAL-1. DMRs around Transcription start sites (TSSs) of the two isoforms showed extensive differentiation between ESCC and normal samples. (d) ESCCAL-1 was significantly differentially expressed and highly abundant in ESCC samples (p-value = 0.0013, FDR=0.0044 by one-way ANOVA). (e) Methylation status of the ESCCAL-1 promoter region was verified on an independent 32 matched normal and ESCC tumor samples using a methylation-specific PCR (MS-PCR) assay; four representative PCR results are shown. M: PCR with methylation primers, U: PCR with unmethylation primers. (f) Quantification of MS-PCR results in the 32 paired normal and tumor samples. Chi square analysis tested for significance between groups. p-value < 0.01. (g) ESCCAL-1 expression is significantly higher in ESCC tumors in an independent cohort of 73 matched normal and tumor samples. *** p-value < 0.005

**Figure 7.**
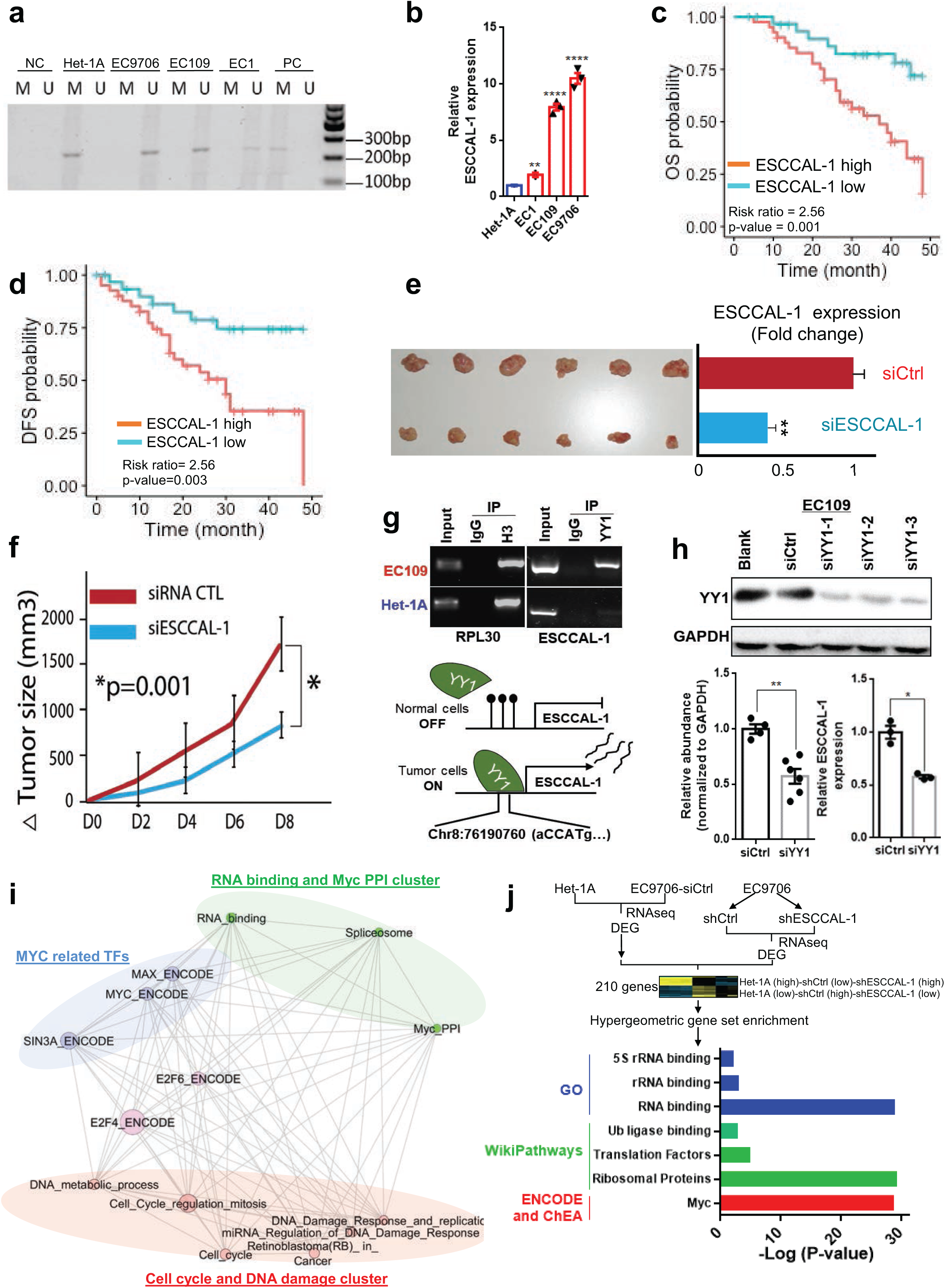
Oncogenic functions of ESCCAL-1 in ESCC. (a) Hypomethylation at ESCCAL-1 promoter regions was confirmed in three different ESCC cell lines using methylation specific PCR. M: PCR with methylation primers, U: PCR with unmethylation primers. Het-1A: an immortalized esophageal epithelial cell line. EC1, EC109 and EC9706: patient-derived ESCC cell lines. (b) ESCCAL-1 expression was significantly higher in three ESCC cell lines (EC1, EC109, and EC 9706) relative to a normal esophageal epithelial cell line (Het-1A). p-value < 0.05. (c-d) ESCC patients with higher expression of ESCCAL-1 exhibit worse overall (OS) and disease-free survival time (DFS), risk ratio=2.56. p-value < 0.005. (e-f) shRNA knockdown of ESCCAL-1 inhibited tumor growth in a tumor xenograft mouse model (N=6, p-value = 0.001). (g) Chromatin immunoprecipitation by YY1 transcription factor protein-directed antibody followed by a standard Polymerase Chain Reaction (PCR) assay in ESCC cancer cells (EC109) and normal esophageal cells (Het-1A). IgG was used as negative control. H3 was used as positive control. (h) Three independent siRNAs targeting various transcript regions of YY1 were transfected into ESCC cell line, 72 hours post-transfection, the total cell lysates were subjected to Western blot assay. Total RNAs were extracted to measure ESCCAL-1 expression using RT-PCR. (i) Nested Gene Set Enrichment Analysis (GSEA) network for ESCCAL-1 gene regulatory network module. Hypergeometric GSEA for gene regulatory network modules (Supplementary Figure 30) was conducted using multiple curated databases (BIOGRID, ENCODE, KEGG, Reactome, GO molecular function). The gene sets are represented as nodes and node size represents the significance of the hypergeometric analysis. Gene sets were further subjected to a hypergeometric test for overlap among them and edges represent significance of overlap; higher the significance for the overlap is indicated by node proximity. 2D projection of 3D network clustering using edge weights were performed using edge-directed spring embedded network layout in Cytoscape 3.2. Ellipses were drawn manually for visualization to demarcate the clusters and nodes were further color coded manually based on their molecular properties. Overexpression of ESCCAL-1 correlated with dysregulation of the cell cycle, DNA repair, RNA binding processing and Myc pathway activation. (j) RNAseq was performed in duplicate in the normal esophageal cell line Het-1A, ESCC cancer cells EC9706 with control siRNA, or EC9706 with a siRNA against ESCCAL-1. Unsupervised hierarchical clustering of differential gene clusters between the three conditions is shown. Differentially expressed genes were selected based on an iterative clustering approach selecting for genes with the top 5% of the most variable and differential gene expression. 210 genes were identified and subjected to GSEA analysis using hypergeometric test in multiple databases as in Figure 4a.

To identify a possible mechanism of ESCCAL-1 upregulation, we examined sequence motifs of known TFs in the ESCCAL-1 hypomethylated promoter region and found a predicted binding site for YY1 from the ENCODE project. YY1 is a TF belonging to the GLI-Kruppel class of zinc finger proteins and contributes to tumorigenesis ^43^. Using ChIP-PCR, we validated YY1 binding at the hypomethylated promoter region of ESCCAL-1 (Fig. 7g). siRNA knockdown of YY1 expression led to decreased expression of ESCCAL-1 (Fig. 7h), indicating YY1 transcriptionally regulates ESCCAL-1 in ESCC.

Since the downstream mechanism of ESCCAL-1’s contribution to ESCC pathogenesis is not clear, we performed a “guilty-by-association” co-expression analysis using RNAseq from ten pairs of normal and tumor samples. The ESCCAL-1 related gene expression modules are enriched in cell cycle pathways, RNA binding and the Myc pathway (Fig. 7i and Supplementary Fig. 29a, b). In order to explore the causative roles of ESCCAL-1 in ESCC progression, we conducted RNAseq in EC9706 cells with ESCCAL-1 knockdown by shRNA (or with shControl). We hypothesized that depletion of ESCCAL-1 could reverse the phenotype of cells alike to a normal cell state, thus we also use RNAseq from a normal cell line, Het-1A. We identified the 210 significantly differentially expressed genes (DEGs) between shControl expressed EC9706 and shESCCAL-1 expressed EC9706 and Het-1A (normal) (gene list in Supplementary table 4) using an iterative clustering approach ^44^. Gene enrichment analysis on these identified DEGs exhibit an enrichment of “RNA binding”, “ribosomal proteins”, and Myc target gene sets (Fig. 7j, Supplementary Fig. 30). These results indicate that ESCCAL-1 participates in the biological process of Myc-mediated regulation of genes, which extends current knowledge on the potential role of Myc signaling in ESCC ^45^. Thus, beyond WNT2-mediated WNT pathway activation, other aberrant signaling pathways activated by ESCCAL-1 upregulation also contribute to ESCC tumorigenesis. Therefore, epigenetic dysregulation promotes ESCC through divergent and multi-factorial mechanisms (diagram in Fig. 8).

**Figure 8.**
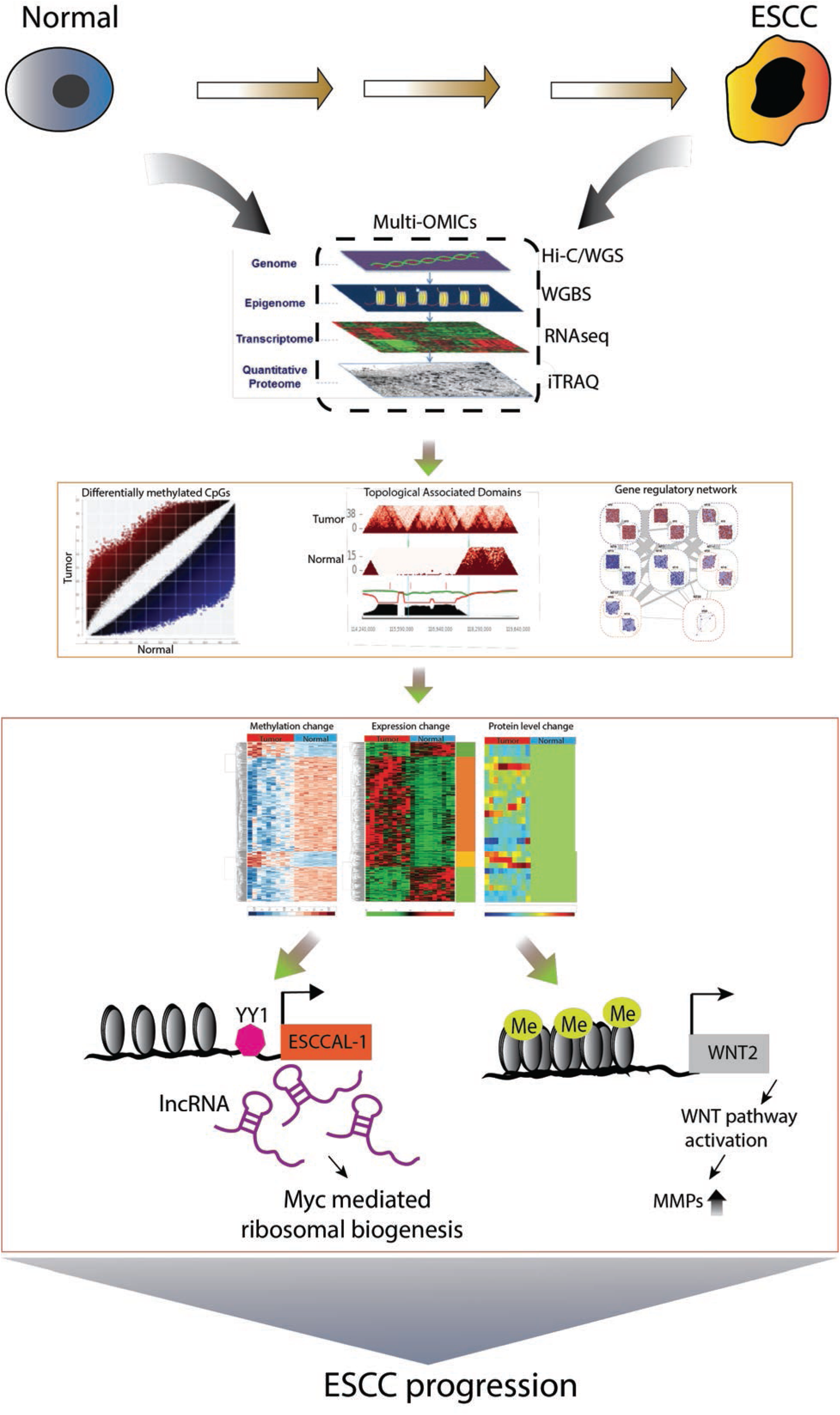
The scheme of multi-omics utilization for defining coding and non-coding driver events in esophageal squamous carcinoma. Each technique for epigenome (genome-wide, high resolution DNA methylome), genomic interaction (interactome), transcriptomic (transcriptome), and proteomic (proteome) analysis define unique alterations during cancer development and progression. The integration of these components in cancer cells with multi-dimensional space measured by orthogonal multi-OMIC analyses informs the understanding of the interplay between genetic and epigenetic dysregulation in ESCC. The datasets facilitate the identification of gene regulatory networks mediated by coding and non-coding transcripts that promote critical tumor phenotypes such as proliferation, invasion, and migration.

## Discussion

The development of ESCC is a complex dynamic biological process that involves multiple steps of genetic and epigenetic alterations. Numerous genetic studies of ESCC at whole genome and exome levels revealed recurrent genetic alterations and related altered pathways such as cell cycle, p53, and AKT/mTOR signaling pathways and Hippo signaling pathways ^4,6,8,46,47^. It remains unclear in ESCC and most other cancer types whether and how the epigenetic landscape contributes to cancer pathogenesis. We performed WGBS, RNA-seq, and proteomic analyses on matched normal and tumor samples along with analysis of TCGA-ESCA datasets and Hi-C sequencing on esophageal normal and tumor cell lines. We observed global hypomethylation (98%) and local hypermethylation across the ESCC genome, consistent with previous studies in colon cancer and other types of cancers ^12,48^. The DMCs alone can discriminate cellular states between tumor and normal conditions, and histological subtypes of esophageal cancer. DNA methylation is a defining feature of cellular identity and is essential for cell development ^49^. Hansen et al. identified cancer-specific differentially methylated regions in colon cancer and showed that such stochastic methylation variations distinguish cancer from normal ^13^ and can be a potential epi-biomarker for early tumor diagnosis or a predictive epi-biomarker for therapeutic outcome. We revealed the heterogeneity of DNA methylation alteration is greater in ESCC relative to normal esophageal tissues and, was a biomarker of inferior clinical outcome. Our findings provide new insight into the potential clinical relevance of epigenetic dysregulation and heterogeneity as a molecular biomarker of clinical outcome in cancer.

We validated a prominently epigenetically altered coding and noncoding gene from the non-canonical cluster (C3) and canonical cluster (C2), respectively, that we defined. The WNT pathway appears to be epigenetically regulated via inactivation of negative regulators (SFRP1/2/4/5 and WIF1) in ESCC ^30^. Our data identified high WNT2 expression in ESCC, along with a highly methylated promoter region. HM450K methylation array and RNA-seq of TCGA ESCA analysis showed high promotor methylation and high gene expression of WNT2. The collective data suggest EZH2-mediated PRC2 repression of WNT2 expression in normal cells. By contrast, we show hypermethylation-mediated de-repression of WNT2 activates the WNT pathway in ESCC. Our data provide new insight into the mechanism of epigenetic dysregulation via non-canonical gene expression regulation in cancer and new insight into the underlying molecular events promoting WNT pathway activation in ESCC.

LncRNA dysregulation is an emerging but poorly understood feature of oncogenesis ^37^. We reported ESCCAL-1 overexpression in ESCC ^20^. Additional studies showed that this lncRNA is overexpressed in multiple types of cancers ^38–40^. Overexpression of this lncRNA promotes cancer cell growth ^50^, invasion^51^ and metastasis ^52^. Here, we discovered that methylation loss of its promoter is a principle molecular mechanism of ESCCAL-1 dysregulation in ESCC. We show that ESCCAL-1 has an epigenetically-mediated causal role in tumor growth and is a biomarker of worse survival of ESCC patients. Interestingly, overexpression of ESCCAL-1 is also related to other types of cancer and it is correlated to drug resistance in lung cancer ^38^. Whether ESCCAL-1 is similarly regulated via epigenetic mechanisms in other cancer types beyond ESCC remains to be investigated in future studies. Nevertheless, suppressing ESCCAL-1 expression, potentially using anti-sense RNA ^53^ or CRISPR-based strategies ^54^, may be a promising therapeutic approach in ESCC and other cancer types.

Our study provides a rationale and a roadmap for delineating the landscape and functional roles of epigenetic dysregulation in cancer at genome-wide high resolution. Further analysis will be required to fully understand the impact of epigenetic dysregulation and heterogeneity on various cancer-associated phenotypes and treatment responses ^14^. Multi-regional WGBS or single cell DNA bisulfite sequencing could facilitate addressing this opportunity in the future.

## Acknowledgements

This study was supported by the National Natural Science Foundation of China (Grants 81171992, 31570899), the Natural Science Foundation of Henan (Grants 182102310328, 162300410279, 182300410374, 192102310096); the Education Department of Henan Province(18B310022,19A320037). This work used the Genome Sequencing Service Center by Stanford Center for Genomics and Personalized Medicine Sequencing Center, supported by the grant award NIH S10OD020141. T.G.B acknowledges funding support from NIH / NCI U01CA217882, NIH / NCI U54CA224081, NIH / NCI R01CA204302, NIH / NCI R01CA211052, NIH / NCI R01CA169338, and the Pew-Stewart Foundations.

## Author contributions

W.C and W.W conceived the project, W.W, H.L, A.Z, S.M performed WGBS, RNA-seq and Proteomic data analysis and construct the figures and manuscript. J.C and N.S.A helped WGBS, multi-omics analyses. G.S performed simulation, A.H.S conducted SNV analysis from RNA-seq, Q.H.X, Y.B.C, J.L, H.G.G, P.J.L, X.Y.S, L.S, P.L.H, and Y.N.L performed experiments. J.W.W and M.Y collected, processed specimen as well as performed some experiments. H.G.X, W.X.T, J.C, J.C.G, and Y.C.G performed Whole Genome Sequencing (WGS), Whole Genome Bisulfite Sequencing (WGBS), Whole Transcriptome Sequencing (RNA-seq), Isobaric Tag for Relative and Absolute Quantitation (iTRAQ) and Hi-C sequencing. F.G.B, E.C helped to discuss the results, T.G.B and M.S guided, discussed and edited the manuscript.

## Competing interests

T.G.B is an advisor to Revolution Medicine, Novartis, Astrazeneca, Takeda, Springworks, Jazz, Array Biopharma and receives research funding from Revolution Medicine and Novaritis.

## METHODS

### Primary esophageal squamous carcinoma (ESCC) specimens

Matched clinical samples of esophageal squamous carcinoma and adjacent normal esophageal tissue are obtained from ten patients (Linzhou Cancer Hospital) as fresh frozen specimens at Translational Medical Center, Zhengzhou Central Hospital, affiliated to Zhengzhou University, China (Supplementary Fig. 1). All samples were surgical resections. The collection of human samples and the protocols for the investigations were under the approval of the Institutional Ethics Committee of Zhengzhou, Henan Province. These specimens were used for sequencing and different assays; WGBS (n=20), WGS (n=6), RNA-seq (n=20) and iTRAQ proteomic assay (n=20). For validation, independent 73 matched ESCC and adjacent normal samples with clinical follow-up were prepared for gene expression analysis.

### Cell cultures

Human ESCC cell lines (EC-109, EC-9706, EC-1) and immortalized esophageal epithelial cell line Het-1A were purchased from the Shanghai Institutes for Biological Science (Shanghai, China). All cell lines were cultured in DMEM medium supplemented with 10% fetal bovine serum (Hyclone, Logan, UT, USA) and maintained at 37°C in a humidified 5% CO_2_ incubator.

### Cell transfection

ESCCAL-1 siRNA, WNT2 siRNA, YY1 siRNA, and their control siRNAs, were synthesized from GenePharma (Shanghai, China). Transfection for EC-109 and EC-9706 cells were previously described ^1^ using Lipofectamine™ 2000 (Invitrogen, Carlsbad, CA, USA) according to manufacturer’s instructions. Transfection efficacy was validated by RTq-PCR.

### Transwell Migration/Invasion Assay

Cell migration and invasion assays were carried out in transwell chambers (Costar, Lowell, MA, U.S.) inserted into 24-well plates. For invasion assay, upper chamber of the transwell plate was coated with matrigel and allowed to solidify at 37 °C and 5% CO_2_ incubator for 30 min. The transfected EC109 and EC9706 cells were trypsinized, resuspended in serum-free culture medium and adjusted to 2×10^6^ cells/ml, added 200 μl cell suspension and 500 μl DMEM containing 10% FBS to the lower chamber. After incubation for 48 h, the transwell chamber was fixed with 10% methanol, followed by the staining with crystal violet. Cells were counted under an inverted microscope. The protocol used for migration assay was similar to the invasion assay without the need to coat the upper chamber of the transwell with matrigel. Each experiment was conducted in triplicate.

### Mouse xenograft experiment

Six-week old male BALB/c immunodeficient mice were purchased from the Shanghai Experimental Animal Center, Chinese Academy of Sciences (Shanghai, China). Animal experimental procedures were carried out according to the Ethical Committee of Zhengzhou University. Mice were housed under a 12 h light/dark cycle and automatically given food and water. The EC9706 cells expressing with ESCCAL-1-siRNA or siControl were subcutaneously injected into back flank of mice as the Knock-down group (*n =* 6) or the control group (*n =* 6). The tumor volumes were calculated as length × width^2^ × 0.5 from day 13 to day 23 every two days, the mice were sacrificed at day 23 after injection.

### Whole genome sequencing (WGS) of three matched tumor and normal tissues

DNA was extracted using the QIAamp DNA Mini Kit (QIAGEN), fragmented using Bioruptor^®^ Pico. Libraries were constructed using VAHTSTM Universal DNA Library Prep Kit for Illumina V3 ND607-02 (Vazyme, Nanjing). Libraries were sequenced with an Illumina HiSeq 4000 to obtain 150 base pair paired-end reads. Base calling was performed with the Illumina Real Time Analysis version 2.7.7 and the output was demultiplexed and converted to FastQ format with the Illumina Bcl2fastq v2.19.0.316. The FastQC package (http://www.bioinformatics.babraham.ac.uk/projects/fastqc) was used to check the quality of the sequencing reads. Sequencing adapters were trimmed from raw reads with Trimmomatic ^2^. We performed mapping, marking duplicates marking, and mutation calling with bcbio-nextgen ^3^, a community developed platform for variant calling. Specifically, Reads were mapped to the human reference genome assembly GRCh37 using BWA-MEM in the BWA package (version 0.7.17) with the default parameters ^4,5^. Duplicates were marked using the tool Sambamba version 0.6.6 ^6^. Somatic mutations include single nucleotide variants (SNVs), small insertions and/or deletions (indels), and structural variants (SVs). The detection of somatic mutations was performed using tumor and matched normal whole genome BAM files generated in the steps described above. We used a series of software packages including VarDict ^7^, MuTect2 ^8^ and Strelka2 ^9^ to detect somatic SNVs and indels, and packages including LUMPY ^10^, Manta ^11^, CNVkit ^12^ and MetaSV ^13^ to detect SVs.

### Whole Genome Bisulfite sequencing (WGBS)

Genomic DNA was extracted with QIAmp DNA Mini kits (Qiagen) from fresh frozen tissue samples. 1μg gDNA was fragmented by sonication with a base pair peak of 300 bp for the resulting fragment, and adaptors were then ligated to both ends of the fragments. Bisulfite conversion was performed to whole genomes of ten pairs of ESCC and matched normal tissues, where converts cytosine residues of the dinucleotide CpG to uracil but leaves methylated cytosine unaffected ^14^. PCR amplification and purification were carried out. The uracil-binding pocket of KAPA HiFi DNA Polymerase has been inactivated, enabling amplification of uracil-containing DNA. The high quality of the library was estimated by The Qubit® 3.0 Fluorometer. Bisulfite conversion success ratio is 99.18% and 99.49%, respectively in normal and ESCC samples (Supplementary Tables 2-3). The WGBS library was sequenced on an Illumina HiSeq2500 sequencers and generated 400M of paired-end reads (2×125bp).

### Whole transcriptome sequencing (RNA-seq)

The RNA was extracted with TRIzol from ten pairs of fresh frozen tissue samples. The input material for total RNA-seq library preparation was 2 μg per sample. Sequencing libraries were generated using NEBNext^®^ Ultra™ RNA Library Prep Kit for Illumina^®^ (#E7530L, NEB, USA) following the manufacturer’s recommendations. Index codes were added to attribute sequences to each sample. Briefly, mRNA was purified from total RNA using poly-T oligo-attached magnetic beads. Fragmentation was carried out using divalent cations under elevated temperature in NEBNext RNA First Strand Synthesis Reaction Buffer (5X). First strand cDNA was synthesized using random hexamer primer and RNase H. Second strand cDNA synthesis was subsequently performed using buffer, dNTPs, DNA polymerase I and RNase H. The library fragments were purified with QiaQuick PCR kits and elution with EB buffer. Terminal repair, A-tailing, and adapter ligation were implemented. The aimed products were retrieved by agarose gel electrophoresis followed by PCR, then the library was completed. RNA concentration of library was measured using Qubit® RNA Assay Kit in Qubit® 3.0 to preliminary quantify and then diluted to 1ng/µl. Insert size was assessed using the Agilent Bioanalyzer 2100 system (Agilent Technologies, CA, USA), and qualified insert size was accurately quantified using StepOnePlus™ Real-Time PCR System (Library valid concentration>10 nM). The clustering of the index-coded samples was performed on a cBot cluster generation system using HiSeq PE Cluster Kit v4-cBot-HS (Illumina) according to the manufacturer’s instructions. After cluster generation, the libraries were sequenced on an Illumina Hiseq 4000 platform to 2×150 bp paired-end reads.

### Proteomic assay and data analysis (Isobaric tag for relative and absolute quantitation, iTRAQ)

Reduced and tryptic digested peptides from samples were labeled with 8 isoberic iTRAQ reagent for an individual run and mixed at an equimolar ratio. Resuspended labeled peptides were pH optimized and separated through strong cation ion chromatography. Samples prepared as such were run through reverse phase LC-MS. The isobaric labeling and LC-MS quantifications were operated in Beijing Genomics Institute (BGI) using optimized quantitative MS-MS protocols ^15^.

For Protein identification and data analysis, IQuant was used^16^. For improved protein identification, a Mascot Percolator and Mascot Parser, a customized post-processing tool was used. The signal to noise ratio was decreased by variance stabilization normalization (VSN). Due to the low abundance or low ionization of peptides, missing the reporter ions is a common phenomenon in isobaric data, and may hinder downstream analysis. A missing reporter was imputed as the lowest observed values to avoid estimation bias. Nonunique peptides and outlier peptide ratios are removed before quantitative calculation ^17^. The weight approach proposed is employed to evaluate the ratios of protein quantity based on reporter ion intensities ^18^. The ratio between normal and tumor samples were generated for each match control pairs. This way three distinct datasets were generated for ten pairs of tumor and normal samples. Sample number 7 (for both tumor and normal, T7 versus N7) was run in each time to standardize among three datasets. For dataset integration, each dataset was normalized by the T7/N7 ratio for the abundance of the protein in all datasets. Dataset normalized as such was represented as a matrix so that protein abundance (row-wise) can be compared across ten different samples as tumor versus normal quantitative ratios (column-wise) generated from three separate runs.

### Hi-C sequencing

The fixed cells (Het-1A, EC109) were resuspended in 1ml of lysis buffer (10 mM Tris-HCl pH 8.0, 10 mM NaCl, 0.2% Igepal CA-630, 1/10 vol. of proteinase inhibitor cocktail (Sigma)), and then incubated on ice for 20 minutes. Nuclei were pelleted by centrifugation at 4 °C, 600x g for 5 minutes, and then washed with 1 ml of the lysis buffer, followed by another centrifugation under similar conditions. After washing twice with restriction enzyme buffer, the nuclei were resuspended in 400μl of restriction enzyme buffer and transferred to a safe-lock tube. Next, the chromatin is solubilized with dilute SDS and incubation at 65 °C for 10 min. After Quenching the SDS by Triton X-100 overnight digestion was applied with 4 cutter restriction enzyme (400 units MboI) at 37°C on the rocking platform. The next steps are Hi-C specific, including marking the DNA ends with biotin-14-dCTP and performing blunt-end ligation of crosslinked fragments. The proximal chromatin DNA was re-ligated by ligation enzyme. The nuclear complexes were reversely crosslinked by incubating with proteinase K at 65°C. DNA was purified by phenol-chloroform extraction. Biotin-C was removed from non-ligated fragment ends using T4 DNA polymerase. Fragments were sheared to a size of 200-600 base pairs by sonication. The fragment ends were repaired by the mixture of T4 DNA polymerase, T4 polynucleotide kinase and Klenow DNA polymerase. Biotin-labeled Hi-C sample was specifically enriched using streptavidin C1 magnetic beads. The fragment ends were adding A-tailing by Klenow(exo-) and then adding Illumina paired-end sequencing adapter by ligation mix. At last, the Hi-C libraries were amplified by 12-14 cycles PCR, and sequenced in Illumina HiSeq platform. Sequencing interacting pattern was obtained by Illumina HiSeq instrument with 2×50bp reads.

### Hi-C data analysis

HiC-Pro algorithm (V2.7.8) was used to process Hi-C data from raw reads to normalized contact maps ^19^. Briefly, (1) valid paired reads were used for the analysis of inter-(Cis) and intra-(Trans) chromosomal interactions. Chromatin interaction frequencies (Ifs) were used to measure the strength of interaction ^20^. The observed/expected number of contacts between all pairs of whole chromosomes were normalized. The interaction between chromosomes was blotted with heatmap (log_10_(valid reads pair) or Circos plot (bin of strongest 1000 pairs). (2) For compartments analysis, Iced+Observed/Expected normalization was used ^21^. Normalized bin pairs were computed Pearson correlation, then the first eigenvector was used to define A/B compartments. (3) Topologically associated domains (TADs). The Insulation method ^22^was used to define the TAD boundary. The TAD heatmap shows at 40kb scale.

### Chromatin Immunoprecipitation (ChIP) with massively parallel DNA sequencing (ChIP-seq)

ChIP assays were performed according to the protocol supplied with the kit (catalog no. 9003) from Cell Signaling Technology. Briefly, EC109 and Het-1A cells were cross-linked with 37% formaldehyde at a final concentration of 1% at room temperature for 10 min. Fragmented chromatin was treated with nuclease and subjected to sonication. Chromatin immunoprecipitation was performed with anti-KMT6/EZH2 antibody (ab195409, Abcam),Anti-Histone H3 (acetyl K27) antibody ChIP Grade (ab4729,Abcam), Anti-YY1 antibody (ab38422,Abcam), rabbit anti-histone H3 (a technical positive control; 1:50) (catalog no. 4620; Cell Signaling Technologies), and normal rabbit IgG (a negative control; 5 μg) (catalog no. 2729; Cell Signaling Technologies). After reverse cross-linking and DNA purification, ChIP-Seq libraries were prepared and sequenced on a HiSeq 4000 sequencer (Illumina, San Diego, CA). To ensure the accuracy of subsequent bioinformatics analysis, the original sequencing data was filtered to obtain high-quality sequencing data (clean data). Quality control of the sequencing data was performed using Sickle (https://github.com/najoshi/sickle) and SeqPrep (https://github.com/jstjohn/SeqPrep). The sequencing output raw reads were trimmed by stripping the adaptor sequences and ambiguous nucleotides and reads with quality scores less than 20 and lengths below 20 bp were removed. The cleaned reads were aligned to human reference genome hg19 using BWA. MACS2 (model-based analysis of ChIP-seq) algorithm^23^ was used for peak calling. The reads of EZH2 binding on WNT2 promoter region were visualized using Integrative Genomics Viewer (IGV, Broad Institute).

### WGBS Data preprocessing

Bisulfite-treated DNAs are further sequenced using Illumina HiSeq 2500 system. Approximately 400M of paired-end reads (125bpx2) and 100 Gbp per sample are generated except one sample (N15). The reads are sent out to the pipeline. Our pipeline consists of four steps (Supplementary Fig. 2a). Firstly, we trimmed using Trim Galore! (v0.4.1) to remove Illumina adaptors with the options of ‘ --paired --length 50 --clip_R1 6 --clip_R2 6 ‘. After trimming, around 90 Gbp of data remained per sample. Then trimmed reads were aligned to the HG19 reference genome using BSMAP (v2.89) with the option of ‘-p 8 -R’. Then SAMtools (v. 1.3.1) was used to sort by genomic coordination and make a bam file index. Picard Tools (v.1.92) is used to remove PCR duplicates. After deduplication, ∼70Gbp remained per sample. Lastly, we ran MOABS (v. 1.3.4) to compute the methylation ratio per CpG with the option of ‘-- cytosineMinScore 20 --skipRandomChrom 1 -p 4 --keepTemp 0 --processPEOverlapSeq 1 -- requiredFlag 2 --excludedFlag 256 --minFragSize 110 --reportCpX G --qualityScoreBase 0 -- trimRRBSEndRepairSeq 0 --trimWGBSEndRepairPE1Seq 5 --trimWGBSEndRepairPE2Seq 5’. Around 95% of CpGs are covered by at least one read.

### WGBS Data Matrix

Given the methylation ratio and coordination computed, we built a data matrix, whose columns are samples and rows are CpGs. The normal tissue WGBS of sample 15 (N15) has generated a small amount of volume (∼4.5Gbp), which is just 6% compared to average (72Gbp) and covers the half of CpGs with notably low coverage (1.58X) than other samples (∼15X). Thus, we decided to eliminate N15 for the further downstream analysis. As a result, we have ten ESCC samples and nine normal esophageal tissue samples. For more robust analysis, we applied the minimum threshold 5X and also selected CpGs that all samples have its methylation ratio. This screening process gave 13 M of CpGs with confident methylation ratio (Supplementary Table 6-7).

### Data quality assurance by TCGA ESCA

To assure our WGBS data (N=19) quality, we aligned our data with TCGA ESCA data (N=202). The methylation data of TCGA ESCA exploited Illumina Infinium HumanMethylation450 Beadchip to measure methylation level for 202 samples; 186 esophageal cancer tissue samples, and 16 esophageal normal tissue samples. To compare TCGA ESCA HM450 methylation ratio and WGBS methylation ratio of our data, we computed the mean methylation ratio of the tumor and normal samples per CpG for both TCGA and our WGBS data. Around 300K of CpGs are in the intersection of TCGA ESCA HM450 and WGBS. We computed the Pearson correlation coefficient (PCC) to measure the representative power in our data set albeit a rather small sample size. First, we calculated the PCC of the tumor and normal from TCGA ESCA HM450 and WGBS. The highest PCC (=0.9674) is between TCGA ESCA normal and WGBS normal since normal tissues are relatively homologous. The PCC (0.9639) between TCGA ESCA tumor and WGBS tumor was followed due to tumor heterogeneity but still showed a high correlation. The third and fourth PCCs are between TCGA ESCA tumor and WGBS normal (=0.9512), and TCGA normal and WGBS tumor (=0.9468) due to the difference between tumor and normal esophageal tissues.

We further assessed how sensitively WGBS could separate ESCA subtypes; ESCC (esophagus squamous cell carcinoma) and EAC (esophagus adenocarcinoma) despite its small sample size and low coverage. We downloaded clinical annotation to match the histological type of each sample. All of the WGBS data is ESCC. The PCC (0.757) between TCGA ESCC and WGBS ESCC is higher than the PCC (0.5554) between TCGA EAC and WGBS ESCC. The PCC between TCGA normal samples and WGBS normal samples is higher than those of tumor samples because of tumor heterogeneity. Overall our WGBS data is as informative as TCGA ESCA HM450 data regarding ∼500K CpG loci and possibly can hold more information about other CpG loci since it covers more CpGs up to 27M.

### Data processing of RNA-seq

RNA-Seq reads were mapped to the HG19 reference genome using STAR (Spliced Transcripts Align to a Reference, v2.4.2a). The expression level of transcript per million (TPM) reads were quantified using RNA-Seq by Expectation-Maximization algorithm (RSEM v1.2.29). The quantified gene expressions of 26,334 transcripts (including coding genes and non-coding genes) were processed in Rstudio console with R programme (v 3.4). Differentially expressed genes between tumor and normal samples were identified using the EdgeR algorithm.

### The algorithm to get DMC (Differentially Methylated CpG)

Among 13M of CpGs, we computed the F-statistics from one-way analysis of variance (ANOVA) to identify confident CpGs. Almost half of them has very low p-values (<0.05). The p-values are further adjusted by Benjamini-Hochberg procedure to compute FDR. 5 M of CpGs has q-values less than 0.05. The 5 M of DMCs is used for the downstream study in this paper. Among 5M of DMCs 97.29% are hypomethylated in ESCC samples while only 2.71% are hypermethylated in ESCC samples (Supplementary Table 8-12).

### Entropy Analysis

Entropy is computed per CpG in both ESCC and normal esophageal cohorts separately as a measure of variance. The ‘entropy’ function was used in the ‘stats’ package of SciPy (v 0.19.1) on top of python3 (v 3.5.2). The bin size was 10%. The distribution of CpG entropy was plotted using MatLab (v. 9.2) ‘plot’ and ‘histogram’ function with the default option.

### DMC enrichment analysis with genomic annotation

We annotated the regulatory elements of the confident DMCs. The genomic coordination of exons an introns were downloaded from UCSC Genome Browser (http://hgdownload.soe.ucsc.edu/goldenPath/hg19/database/refGene.txt.gz). The promoter is computed 4.5kbp upstream and 500bp downstream given TSS. The genomic coordinates of enhancers are downloaded from VISTA Enhancer Browser (https://enhancer.lbl.gov). We learned that hypomethylated DMCs outnumbers hypermethylated DMCs in ESCC, 4.95M vs. 138K, respectively. Among hypermethylated in DMCs in ESCC, 83.67% has overlapped with regulatory elements while only 56.77% of hypomethylated DMCs has overlaps. Such overlaps were further dissected into enhancers, promoters, exons, and introns (Supplementary Fig. 4).

### DMC enrichment analysis with functional annotation

We performed functional annotation on the confident DMCs except for protein-coding RNAs. Functional annotation includes long noncoding RNA (lncRNA), antisense RNA, Micro RNA (miRNA), small nuclear RNA (snRNA), small nucleolar RNA (snoRNA), small cytoplasmic RNA (scRNA), ribosomal RNA (rRNA), vaultRNA and Mt_tRNA. The functional annotation was downloaded from the GENCODE project (https://www.gencodegenes.org/releases/27lift37.html). The composition of the functional annotation is illustrated. (Supplementary Fig. 5). We deconvoluted functional mapping of hypermethylated DMCs and hypomethylated DMCs. The majority of the hypermethylated DMCs are mapped to antisense RNAs (58.01%) followed by lncRNA (39.28%) while that of the hypomethylated DMCs are mapped to lncRNAs (63.08%) followed by antisense RNA (29.89%).

### Transcription factor binding site analysis

Transcription factors and their binding sites annotation were downloaded from the ENCODE project (http://hgdownload.cse.ucsc.edu/goldenpath/hg19/encodeDCC/wgEncodeRegTfbsClustered/). First of all, we computed the proportion of each transcription factor. With regard to base-pair counting, POLR2A the largest contributor to the composition, followed by CTCF (Supplementary Fig. 8a). We mapped DMCs to transcription factor binding sites and calculated the composition of transcription factors both in DMCs and HG19. Enrichment was computed as a ratio of the proportion of TFBS mapped to DMCs over TFs in HG19 (Supplementary Fig. 8b). Relatively DMC are most enriched in SUZ12 followed by EZH2, which are the component of the Polycomb Repressive Complex 2 (PRC2) (Supplementary Fig. 8c). Methylation type has been studies among top 20 TFs that DMCs are enriched. Hypomethylated DMCs dominated in the most TFs except for three TFs; SUZ12, EZH2 and CTBP2, which is related to endometrial cancer pathway and WNT pathway (Supplementary Fig. 8d).

### TSS Methylation level analysis

The methylation ratio was summarized in every 200-bp window relative to transcription start site (TSS). Then the methylation ratio per bin was normalized and averaged for both ESCC and normal cohorts. Methylation conversion is observed both upstream and downstream given TSS. The normalized methylation ratio of ESCC samples is higher between around −3000bp to 7000bp given TSS. The graph was made by R with spline interpolation with default options.

### Differentially Methylated Region (DMR) calls and landscape in whole genome

DMR was computed from 5Ms of confident DMCs. The window size is flexible as long as any two CpGs locate in 150bp and have consistent methylation pattern either keeping hypermethylated or hypomethylated. The criteria make sure the minimum CpG density is at least 0.01. DMRs peak size is of 150-350bp and CpG density peak is of 0.04-0.05 (Supplementary Fig. 7a, b). The overall distribution of DMRs in HG19, genomic annotation and functional annotation is illustrated in Supplementary Fig. 7c, 7d.

### Integrating and clustering methylome and transcriptome data

To integrate methylomic and transcriptomic data, we focused on methylation in the promoter regions which are defined 4.5Kbp upstream and 500bp downstream given TSS. We aligned DMRs that are identified previously with *q* value < 0.01 to the promoter regions of all genes. We also increased the promoter size up to 11 Kbps to see if any level of extension of promoter size can affect the final gene set whose promoters hold any DMRs, but that added only four more genes. As a result, we learned that 5085 genes have significant DMRs in their promoter regions. High-throughput transcriptome sequencing (RNA-Seq) was conducted. Around 20,000 gene expression levels are estimated for both normal and ESCC pairs of ten patients. Significantly differentially expressed transcriptomes between two cohorts are selected given *p* value < 0.05. Total 4768 genes are significantly highly expressed in the ESCC cohort or the normal cohort.

The number of genes in the intersection, in other words, genes whose promoters are differentially methylated and expressions are significantly different between normal and ESCC cohorts, are 694 in total. We categorized those genes into four groups; (1) C1: Hypermethylated in their promoters with low gene expression in ESCC (2) C2: Hypomethylated in their promoters with high gene expression in ESCC (3) C3: Hypermethylated in their promoters with high gene expression in ESCC (4) C4: Hypomethylated in their promoters and low gene expression in ESCC. The genes in the former two groups (C1 and C2) are explained by canonical promoter methylation and gene expression model while the genes in the latter two groups (C3 and C4) are not. The gene list of the four groups is in the Supplementary Table 13-16. The heat map with dendrograms is made using ‘clustergram’ command of MatLab v9.4 with parameters of ‘Standardize’, ‘Row’, ‘Colormap’, ‘redgreencmap’.

### CTCF analysis in gene promoter and gene body

The genomic coordination of CTCF was retrieved from ENCODE Project (https://www.gencodegenes.org/releases/27lift37.html). We identified the number of CTCF binding site overlapped with promoter regions by BEDTools (v2.26.0) with ‘intersect -wa -wb -a -b’ option. The CTCF ratio in the gene promoter region is highest in C3 (2.2).

### Gene body methylation level analysis

DMCs (FDR<=0.05) in the gene body and promoter region is selected for C1 and C3 to search hypothesis of noncanonical correlation in C3, where promoters are hypermethylated with high gene expression. Overall gene body is hypomethylated for both C1 and C3, but such reverse methylation between promoter and gene body is much prominent in C1 genes.

### Copy number alteration inferred from RNAseq

CNVkit-RNA ^24^ was used to infer copy number alterations from RNAseq reads. The segments and recurrent copy number gains or loss across samples were generated and plotted using GISTIC 2.0 algorithm (Supplementary Fig.17).

### Simulation of CpG methylation heterogeneity

Simulations of Ornstein-Uhlenbeck processes have been performed for CpG islands at selected promoter regions in both normal and cancer samples ^25^. These simulations model the epigenetic landscape as highly regulated stochastic processes in normal tissues, where methylation levels deviate only minimally around the equilibrium mean (like a ball in a narrow valley). In cancer tissue, the regulation is disrupted, and methylation levels are stochastically driven away from the equilibrium (a ball is in a broader valley). This allows us to understand changes beyond mean differential methylation levels. Hypermethylated and hypomethylated regions can be explained by a change in equilibrium methylation levels together with an increase in stochastic deviation. Additionally, this model helps to understand variation in low-to-low and high-to-high differentially methylated regions. For the data in this study, the model works well for regions with single-peaked or broad methylation distributions. The histogram shows methylation variation by the stochastic simulation if more samples collected (Supplementary Fig. 6d).

### Co-expression analysis of RNAseq

Co-expression analysis was conducted in R environment using RedeR package ^26^. RedeR package has a strong statistical pipeline for several network analysis. Here, we have used co-expression analysis, computing a null distribution via permutation and returning the significant correlation values. We performed 1000 permutations to build the null distribution and used Pearson correlation. We considered correlations with p-value less than 0.01 with FDR adjustment as significant. The hierarchical clustering analysis used the complete method and considers the distances of each individual component to progressively computing the clusters until it finds a stable state. The final result is a dendrogram presenting hierarchical leaves, which has been used to plot the network. To clear the visualization, clusters ware nested using the fourth level of dendrogram to build the nests (Supplementary Fig. 29).

### Methylation-specific polymerase chains reaction (msPCR) for ESCCAL-1 and WNT2 validation

DNA was extracted from cells using the AllPrep DNA mini kit (Qiagen) according to the manufacturer’s instructions and was quantified by NanoDrop analysis. Bisulfite modification was carried out on 200-500 ng of DNA using the EZ DNA Methylation-Gold Kit (Zymo Research) according to the manufacturer’s instructions. Methylation-specific PCR (MS-PCR) analysis at BioRad T100TM Thermal Cycler with 20 μL reaction mixtures. Primers for modified methylated sequences and modified unmethylated sequences were listed below. The PCR reactions were carried out under the following conditions: 95 °C for 10min, 95 °C for 30 s, 48.5(51) °C for 30 s and 72 °C for 30 sec for a total of 45 cycles, 72 °C for 10min. 5ul PCR product was used for electrophoresis on 2% agarose gel, the methylated strip and unmethylated strip were analyzed by gel imaging analyzer.

Representative results showing the ECSSAL-1 promoter methylation status identified by MS-PCR in ESCCs. Lanes M and U indicate the amplified products with primer recognizing methylated and unmethylated sequences, respectively. NC, negative control; PC, positive control.

Primers for msPCR:

WNT2:

Methylated Forward: CGCGCCCGTGCGCGTGGACTTA

Unmethylated Forward: TGTGTTTGTGTGTGTGGATTTA

Methylated Reverse: CGCATGGCGCCCGCACACGGAGT

Unmethylated Reverse: CGTATGGTGTTTGTATATGGAGT

ESCCAL-1:

Methylated sense: TGCGCCAGCCGAAGCAGGGCGA

Unmehylated sense: TATATTAATTAAAATAAAATAA

Methylated antisense: CGAGACTCCGTGGGCGTA

Unmethylated antisense: TAAAATTTTATAAATATA

### Gene enrichment analysis

Curated gene sets were collected from multiple curated databases such as ENCODE, CHEA, EnrichR, KEGG, WikiPathways, Reactome, GO molecular function, Panther and BIOGRID. Overlapping between differentially methylated and/or expressed genes were estimated for significance using hypergeometric statistics. P values were adjusted to Bonferroni correction. −Log_10_(p-value) for significance was measured and compared.

### ChIP-PCR

Chromatin immunoprecipitations were performed using digested chromatin from EC-109 cells or Het-1A cells and the indicated antibody YY1. The antibody Histone H3 (D2B12) as a positive control. Purified DNA was analyzed by standard PCR methods using SimpleChIP® Human RPL30 Exon 3 Primers and ESSCAL-1 primers. equal amounts of total genomic DNA (Input) were used for immunoprecipitation in each condition.

Primers for YY1 ChiP-PCR were:

ESCCAL1-YY1 Forward: TTTGAAATAATGAGTTATGAG

ESCCAL1-YY1-Reverse: AGGGAACTCCCTGACCCCTTGC

### Statistics

The student t-test was used for two group mean comparison. One-way ANOVA was used for multiple groups comparison. Hypergeometric test was used in gene set enrichment analysis. Multiple hypothesis test was adjusted with Benjamini-Hochberg method. Th standard α = 0.05 was used as cutoff, the null hypothesis is rejected when p-value < 0.05. Different significant levels were used: * p-value < 0.05; ** p-value < 0.01; *** p-value < 0.005; **** p-value < 0.001. The 95% confidence interval (CI) for the median duration of PFS and overall survival were computed with the robust nonparametric Brookmeyer and Crowley method. Hazard ratio with 95% CI and p*-*values were calculated with the ‘Cox proportional-hazards regression model with survival’ package in R.

### Computational resources and code sharing for reproducibility

Most of the analysis was done using SCG4 cluster of the Genome Sequencing Service Center by Stanford Center for Genomics and Personalized Medicine Sequencing Center. The SCG4 cluster has 20 compute nodes, each with 384 GB RAM, 56 CPUs each, and 40 compute nodes with 16 and 48 CPUs and 10GbE connectivity. It shares 4+ PB of storage NIH dbGaP compliant and have 350+ software packages installed.

